# A vector system for single and tandem expression of cloned genes and multi-colour fluorescent tagging in *Haloferax volcanii*

**DOI:** 10.1101/2024.03.13.584740

**Authors:** Solenne Ithurbide, Roshali T. de Silva, Hannah J. Brown, Vinaya Shinde, Iain G. Duggin

## Abstract

Archaeal cell biology is an emerging field expected to identify fundamental cellular processes, help resolve the deep evolutionary history of cellular life, and contribute new components and functions in biotechnology and synthetic biology. To facilitate these, we have developed plasmid vectors that allow convenient cloning and production of proteins and fusion proteins with flexible, rigid, or semi-rigid linkers in the model archaeon *Haloferax volcanii*. For protein subcellular localization studies using fluorescent protein (FP) tags, we created vectors incorporating a range of codon-optimized fluorescent proteins for N- or C-terminal tagging, including GFP, mNeonGreen, mCherry, YPet, mTurquoise2 and mScarlet-I. Obtaining functional fusion proteins can be challenging with proteins involved in multiple interactions, mainly due to steric interference. We demonstrated the use of the new vector system to screen for improved function in cytoskeletal protein FP fusions, and identified FtsZ1-FPs that are functional in cell division and CetZ1-FPs that are functional in motility and rod cell development. Both the type of linker and the type of FP influenced the functionality of the resulting fusions. The vector design also facilitates convenient cloning and tandem expression of two genes or fusion genes, controlled by a modified tryptophan-inducible promoter, and we demonstrated its use for dual-colour imaging of tagged proteins in *H. volcanii* cells. These tools should promote further development and applications of archaeal molecular and cellular biology and biotechnology.

## Introduction

Archaeal cell biology is an emerging field in evolutionary and developmental biology. The last ∼15 years has seen several developments that have improved the study of archaea that often grow under conditions such as high temperature that challenge the application of established laboratory techniques, such as cell cycle synchronization and live-cell imaging in the model thermoacidophilic archaeon *Sulfolobus acidocaldarius* [1–3]. Yet some archaea such as the halophilic model archaeon, *Haloferax volcanii*, are well suited to most cell biology studies due to their ease of growth under readily available laboratory conditions [4], powerful genetic and biochemical tools [5, 6], and relatively large discoidal or rod shaped cells that may be visualized clearly by live cell microscopy [7–11].

A proper understanding of protein functions in cell biology usually necessitates imaging the subcellular localization of proteins in live cells. One of the main techniques for studying protein localization in genetically tractable organisms is the expression of a fusion between the protein of interest and a fluorescent protein (FP) for detection by fluorescence microscopy [12]. However, protein tags can interfere with the proper function and localization of the protein of interest, which can complicate interpretation and might lead to false conclusions [13, 14]. Proteins involved in large assemblies have been especially challenging, due to steric interference. For example, GFP localization studies of the bacterial cell division protein FtsZ started in the 1990s [15] and, whilst researchers have improved substantially on the earliest fusions, to our knowledge there has been no fully-functional FtsZ-FP identified, despite extensive testing of combinations of fluorescent proteins at different insertion sites in FtsZ [16, 17]. Nevertheless, studies with non-fully functional FP tagged proteins have led to significant advancements in cell biology, mainly by expressing fusion proteins to supplement rather than fully replace the endogenous untagged protein of interest. This approach often does not noticeably interfere with the process under investigation and provides localization data that is expected to closely resemble that of the native protein [15, 16]. However, it is important to evaluate the unpredictable functionality of FP-tagged proteins with *in vivo* functional assays and aim to find highly functional fusions, particularly when applying quantitative or superresolution techniques aimed at resolving and measuring detail of subcellular assemblies where all the target protein molecules in the cell are to be tagged and visualized [16]. Several strategies have been used to improve the functionality of fusion proteins including the testing of different fluorescent tags, linkers of variable size and flexibility between the fused proteins, or the insertion of the FP at different positions in the protein of interest [17, 18].

In recent years, FPs have been applied to archaea to visualize subcellular structures [10, 11, 19, 20]. However, like in bacteria, the FP-tagged proteins are typically produced at moderate concentrations in the presence of the untagged wild-type (WT) copy to allow normal function of the cellular processes under investigation, where the fluorescently tagged proteins may not fully complement the phenotype of interest in the corresponding gene-deletion background. For example, expression of GFP and mCherry fusions to the *H. volcanii* cytoskeletal proteins FtsZ1 only partially complement the cell division or shape defects of the Δ*ftsZ1* [10, 21]. Furthermore, other fusions like FtsZ2-GFP or GFP-SLG are disruptive to division when expressed in the wild-type background [19, 21]. Such effects are potentially exacerbated when expressing two tagged proteins.

To address these challenges, we have created and tested a set of plasmid vectors for generating N- or C-terminal protein fusions with three different linkers (flexible, semi-rigid and rigid), and we generated sets of plasmids for the ready fusion of genes of interest to a selection of five codon-optimised recent-generation FPs. This system was also designed to allow straightforward generation of tandem (bicistronic) dual-expression plasmids for co-localization, or other co-expression studies with untagged proteins, and we identified improved, highly functional FP fusions to cytoskeletal proteins FtsZ1 and CetZ1 in *H. volcanii*.

## Materials and Methods

### Materials, strains, and growth conditions

Media constituents, peptone, bacteriological agar, and casamino acids, were obtained from Oxoid to facilitate haloarchaeal growth [22] and other compounds were obtained from Sigma-Aldrich [8]. For plasmid construction and subcloning, described below, we used *Escherichia coli* DH5α, and obtained oligonucleotide from Integrated DNA Technologies, enzymes from New England Biolabs, and synthetic fragments and genes encoding the fluorescent proteins from Genscript (Supplementary Table 2).

*H. volcanii* strains were cultivated routinely in Hv-Cab medium [8] supplemented with uracil or hypoxanthine and thymidine where required [23]. Cultures were incubated at 42 or 45°C with shaking (200 rpm). *H. volcanii* transformation was performed as described [24], with plasmids purified from the methylation-deficient *E. coli* C2925 (NEB). Cloned gene expression was induced by inclusion of the indicated concentration of L-tryptophan (Trp) in the growth medium throughout cultivation.

*H. volcanii* strains were based on H26 (ID621) or H98 (ID6) [23], which carried the empty vector pTA962 [5] when used as wild-type controls for in vivo assays. Strains carrying deletions of *ftsZ1* or *ftsZ2* in the H98 background have been previously described (ID76 and ID77, respectively) [21]. A strain carrying an in-frame deletion of *cetZ1* (removing codons 8-346) based on the H26 (Δ*pyrE2*) parent strain was generated. Primers for PCR amplification of the *cetZ1* flanking regions for homologous recombination were as follows, for the upstream flank, 2204_US_F (ccggcc**aagctt**gcgagttcgtctccttcacga) and 2204_IF_US_R (<u>gaattcgccgcccgaagatct</u>tccgatcattgcgagcttc), and for the downstream flank, 2204_IF_DS_F (<u>agatcttcgggcggcgaattc</u>tcgggcgtgacgaacg) and 2204_DS_R (ccccc**ggatcc**acgtctgctcggcttgttgc). The two PCR products were combined and used as the template for overlap extension PCR to join them and create the deletion construct (the underlined bases in the primers above representing the overlap region that would replace the *cetZ1* coding region). The spliced PCR product was cloned at BamHI and HindIII sites (bold text in primers above) in pTA131 [23], and the expected sequence was confirmed by Sanger sequencing (Australian Genome Research Facility). This new plasmid (pTA131_2204_IF) was used to transform *H. volcanii* H26 with the two-step selection procedure (‘pop-in-pop-out’) as described [23]. A transformant designated *H. volcanii* ID181 was selected that showed complete exchange of the wild-type *cetZ1* for the deletion, and CetZ1 was also not detected by western blotting.

### Construction of Vectors for producing fusion proteins and tandem expression

To construct the pHVID vector series (Figure 1), we first made a plasmid that silently removes the two XbaI sites in a vector based on pTA962 [5], as shown in Supplementary Figure 1A, resulting in pIDJL114_noXbaI. To enable tandem gene cloning (Figure 1C), p.*tnaA* was first modified (p.*tnaA*\*) to include an XbaI restriction site at the 3’ end. Alignment of several haloarchaeal species’ p.*tnaA* sequences was used to identify regulatory regions of the promoters to help guide the introduction of XbaI to conserve the regulatory elements and their functions (Supplementary Figure 1B and 1C). Synthetic DNA fragments containing one of the three *p.tnaA\**-linker-MCS sequences (Figure 1A) were then cloned into pIDJL114_noXbaI between ApaI and NotI, thus replacing the p.*tnaA*-FtsZ1-mCherry fragment in pIDJL114_noXbaI to generate the three base vectors, pHVID1-3 (Figure 1A). Genes encoding the six fluorescent proteins (Supplementary Table 2) were then subcloned in-frame into each of pHVID1-3, between EcoRI-NheI to generate sets of plasmids for C-terminal tagging (pHVID4 to pHVID21) and between NdeI-BamHI for N-terminal tagging (pHVID22 to pHVID39) (Supplementary Table 1, Figure 1B). Sequences of all cloned regions generated by PCR were verified by Sanger sequencing. To create tandem expression plasmids, the gene to be placed in the downstream position (distal with respect to *p.tnaA*) was isolated via plasmid digestion with XbaI+NotI (Figure 1C) and then ligated to the appropriate pHVID-based plasmid containing the gene for the proximal position (digested with the compatible NheI+NotI).

**Figure 1:**
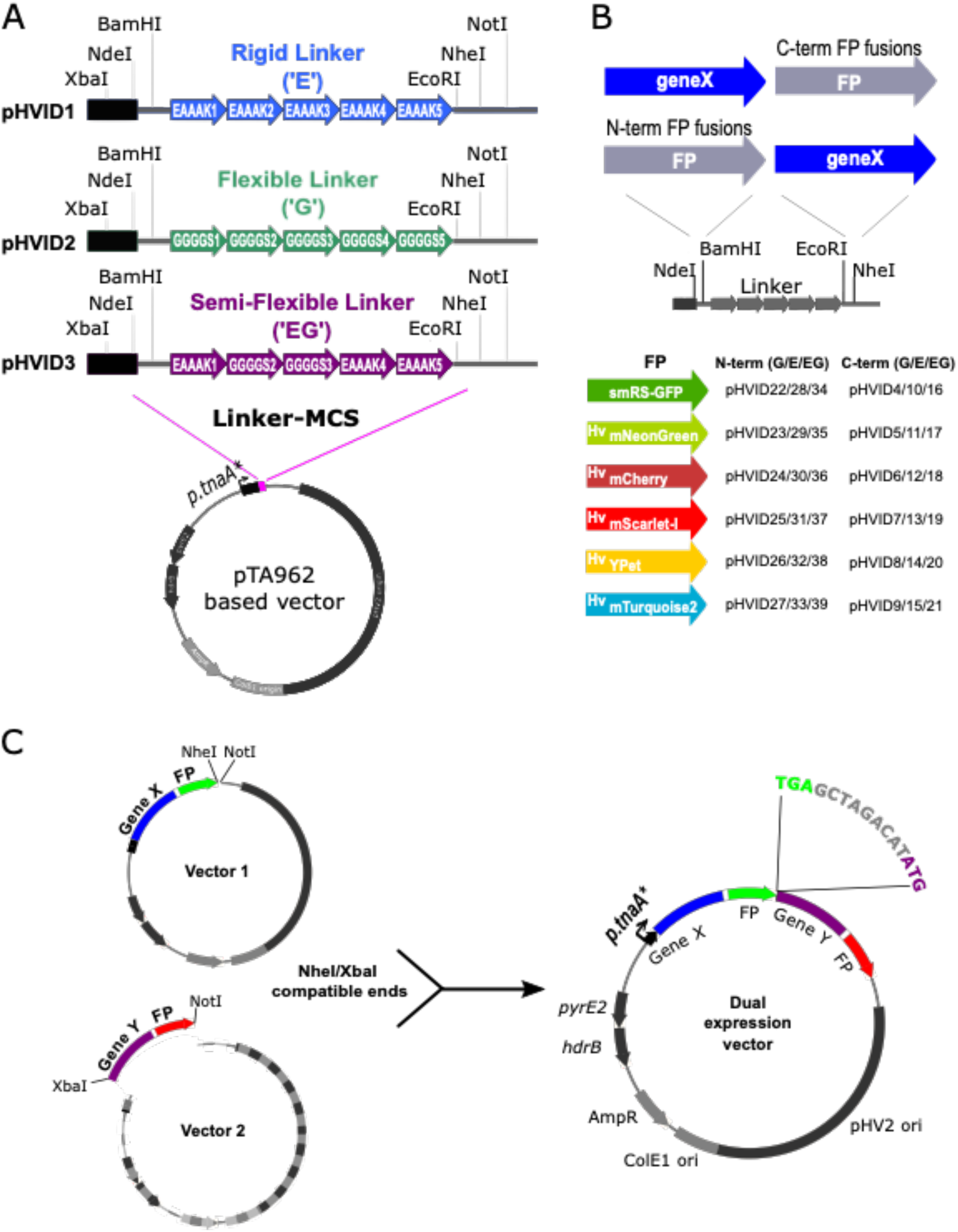
A vector set for construction and production of fusion proteins in *H. volcanii*. (A) The vectors are based on the pTA962 shuttle vector containing the *E. coli* ColE1 origin and AmpR marker gene for propagation in *E. coli*, and the *H. volcanii* pHV2 origin and tandem *pyrE2* and *hrdB* marker genes. Vectors pHVID1-3 contain the three indicated linkers. (B) Outline of construction of N-terminal and C-terminal fusions, and the set of constructed plasmids (pHVID4-39) encoding fluorescent proteins (Hv, codon-optimised) ready to be fused to a gene of interest (‘geneX’). (C) Construction of tandem expression plasmids containing the two fusion genes expressed from the same p.*tnaA* promoter in an operon-like organisation with a distance of 9 bp between them (expanded sequence shown).

### Cell shape and motility assays

To culture *H. volcanii* for measuring rod cell formation during early-log phase strains were first streaked onto an Hv-Cab agar plate and incubated at 45°C for 3-5 days. Single colonies were inoculated into Hv-Cab medium and allowed to grow until an approximate OD600 of 0.6, and then 3 μL was diluted into 10 mL Hv-Cab medium supplemented with 2 mM Trp and allowed to grow for 24 hours before imaging.

Soft agar motility assays of *H. volcanii* strains were carried out using Hv-Cab with 0.3 % (w/v) agar supplemented with 2 mM Trp in six-well culture plates. For each of three replicates, 2 μL of mid-log phase (OD600 ∼0.4) culture was spotted on the surface of the soft agar and allowed to adsorb. Plates were then incubated in a sealed plastic bag with minimal disturbance at 42°C for four days. Images of the motile colonies were obtained using a gel documentation system with white reflected light illumination. The diameter of the ring of motile cells was taken as the mean of two perpendicular measurements, which was then normalized as a percent of the wild-type control *H. volcanii* strain.

### Coulter cytometry

Samples of 100 µL of cells maintained in exponential growth for at least 2 days were diluted (1:100) in 18 % (w/v) Buffered Salt Water (BSW) [8]. The cell volume was analysed with a Multisizer 4 Coulter Counter (Beckman-Coulter). Runs were completed in the volumetric mode (100 μL), with a 30 µm aperture tube, the current set to 600 μA and gain set to 4. Volumetric data were displayed in frequency distributions with 300 bins (from 0.113 µm^3^ to 3054 µm^3^) with a logarithmic bin spacing, taking the central volume value as the bin value and the number of cells was expressed as a percent of the total.

### Fluorescence microscopy and image analysis

For localization studies of FtsZ1 and FtsZ2 fusion proteins, samples of cultures maintained in exponential growth for at least 2 days were mounted on a 1% agarose pad (containing 18% BSW). Image acquisitions were performed on a GE DV Elite microscope equipped with an Olympus 100X UPLSAPO/NA 1.4 objective and a pco.edge 5.5 sCMOS camera. GFP and mNeonGreen fluorescence were acquired with a GFP/FITC filter set (ex: 464-492 nm, em: 500-523 nm), mCherry and mScarlet-I fluorescence with an mCherry filter set (ex: 557-590 nm, em: 598-617 nm), YPet fluorescence with a YFP filter set (ex: 497-527 nm, em: 537-549 nm), and mTurquoise2 fluorescence with a CFP filter set (ex: 400-453 nm, em: 463-487 nm).

Micrograph image analysis was done using ImageJ (Schneider et al. 2012). Cell circularity (4π × area / perimeter^2^) was determined using the MicrobeJ plug-in [25] by taking the interpolated cell contour generated after thresholding using the particle medial axis. To measure total fluorescence intensities of individual cells, the background for each fluorescent channel was first subtracted using a rolling ball size of five pixels. Cells were detected in the phase-contrast channel by generating a ‘Default’ threshold, and the mean and sum of pixel intensities within the cell area (‘Raw Integrated Density’) were measured in each fluorescent channel using the threshold generated on the phase-contrast image as a reference. Finally, the ratio between mCherry and mTurquoise2 intensity for individual cells was calculated.

### Western blotting

Total cell extracts from cells in logarithmic growth were obtained by resuspending cell pellets in reducing sample buffer to a cell density of OD600 = 5 and then incubated at 90°C for 5 min and vortexed. Samples were subjected to SDS-PAGE and then electro-transferred to a nitrocellulose membrane, using standard methods (Bio-Rad). Rabbit primary antibodies have been described previously for *H. volcanii* FtsZ1 (1:1000 serum dilution), FtsZ2 (C-terminal amino acids 383-400) (1:2000 serum dilution) and CetZ1 (1:1000 serum dilution) [10, 21]. The secondary antibody was a Donkey Anti-Rabbit IgG HRP-conjugate (AbCam 16284, diluted 1:5000). Antibody dilutions and wash steps were done using TBST (50 mM Tris-Cl pH 7.4, 150 mM NaCl, 0.05% (v/v) Tween-20). Detection by chemiluminescence was done with the SuperSignal West Pico PLUS reagents (Thermo Fisher Scientific).

## Results

### Plasmid vectors to facilitate construction of fusion proteins and single or dual protein production in H. volcanii

Identifying a functional fluorescent protein (FP) fusion can be challenging and often requires optimization and iterations including testing the position of the FP compared to the protein of interest, the type of FP, and the properties of the linker joining the protein of interest and the FP. To allow for testing several of these parameters, we made a vector set that enables the straightforward generation of fusion proteins for expression in *H. volcanii* with different types of linkers and *H. volcanii* codon-optimised FPs and the additional option of tandem expression of two genes or fusions (Figure 1). The vectors are based on the *E. coli-H. volcanii* shuttle plasmid pTA962, which contains a tryptophan-inducible promoter, *p.tnaA*, for the control of cloned gene expression in *H. volcanii*, and an adjacent multiple cloning site (MCS) [5]. The original *p.tnaA*-MCS region was replaced to create three new base vectors containing each of three different linkers downstream of a *p.tnaA* promoter (Figure 1A and Supplementary Figure 1). Highly flexible linkers can be designed by including repeats of small amino acids that do not form stable secondary structure, ([GGGGS]n) [26, 27], while rigid linkers, aiming to hold the protein of interest and tag further apart, can be designed with repeats of an α-helix forming motif, ([EAAAK]n). Li et al. [28] surveyed combinations of five units of these motifs to create linkers with a range of flexibilities. We selected three 25-amino-acid linkers from this library: a rigid linker (EAAAK5, or “E-linker”), a flexible linker (GSSSS5, or “G-linker”), and a semi-rigid linker (EAAAK1GGGGS2EAAAK2, or “EG-linker”). The resulting vectors allow the insertion of fusion partners N-terminal (NdeI-BamHI sites) and C-terminal (EcoRI-NheI sites) relative to the linker (Figure 1B).

We then cloned the open reading frames (ORF) of six fluorescent proteins individually into the N- and C-terminal positions, to create 12 plasmids for each of the three linker types that are ready for cloning fusion partners in the alternate position (Figure 1B). Four recent-generation fluorescent proteins were selected that were favourable for brightness, short maturation time, low bleaching and amenability to co-localization and FRET-pair studies: mNeonGreen [29], YPet [30], mTurquoise2 [31], and mScarlet-I [32]. Open reading frames (ORFs) encoding the four fluorescent proteins and the mCherry used previously [10] were synthesised and codon-optimized for *H. volcanii*, and the previously used GFP variant was also included (Figure 1B) [10, 33]. This set of plasmids should facilitate the identification and use of functional fluorescent-tagged proteins by generating up to 36 possible combinations of linker, fluorescent protein, and N-/C-terminal position with a protein of interest. The optimal linker and fluorescent protein for tagging the protein of interest may be identified empirically using *in vivo* protein functional assays, such as phenotypic complementation of a strain carrying a knock-out mutation of the gene of interest.

The design of the new *p.tnaA*-MCS in vectors pHVID1-3 also incorporated sites that allow subsequent subcloning to combine two genes in tandem, creating a synthetic operon controlled by the single *p.tnaA* promoter for dual expression studies (*e.g.*, two-colour fluorescence imaging). To enable this, we had modified the original *p.tnaA* to incorporate an XbaI restriction site near its 3’-end (Figure 1A and Supplementary Figure 1, *p.tnaA*\*). With this design, XbaI and NotI can be used to excise one fusion construct for ligation to the compatible ends of another NheI/NotI linearized plasmid to generate the required tandem construct (Figure 1C).

### Screening fluorescent protein fusions to the C-terminus of FtsZ1 for function in H. volcanii

To demonstrate the utilisation of the fluorescent proteins and vectors, we investigated some known cytoskeletal proteins in *H. volcanii*, which form distinct subcellular structures and show readily observable phenotypes when deleted [10, 21]. This facilitates screening for functional fusion proteins via complementation studies, and comparisons of localization patterns with different linkers and FPs. We started by investigating the two FtsZ proteins involved in *H. volcanii* cell division [21]. Although the previous *H. volcanii* FtsZ1-GFP and -mCherry fusions (both containing a 2 amino-acid linker, GS) clearly localized to the midcell division ring, they only partially restored cell size (division) compared to the untagged FtsZ1 when expressed from a plasmid in the Δ*ftsZ1* background [21]. The same study showed that FtsZ2-GFP (with GS linker) localized at mid-cell in the wild-type background but showed no rescue of the cell division defect in the Δ*ftsZ2* background. When used at a low level of supplemental expression in the wild-type background, it could be used to investigate the localization of FtsZ2 at the midcell division site. However, moderately higher expression caused defects in cell division, indicating that the FtsZ2-GFP fusion localises to the division site and acts in a dominant inhibitory manner in cell division. Given these previous findings, we started by using the vector series to make and screen FtsZ1 fusion proteins for improved functionality with the tag placed on the C-terminus of FtsZ1, enabling comparison with the original C-terminal constructs [10, 21].

We cloned FtsZ1 into the proximal position in the sets of plasmids containing the C-terminal fluorescent proteins (*i.e.*, pHVID4 to pHVID21, Figure 1B), generating 18 combinations of FtsZ1, linker and FP fusions. The functionality of these fusions was assessed by measuring the cell volumes in mid-log cultures of *H. volcanii* Δ*ftsZ1* strains carrying the individual plasmids by Coulter cytometry to test for restoration of the cell division defect (larger cell volume) caused by Δ*ftsZ1* (Figure 2A). Secondly, fluorescence microscopy was used observe the subcellular localization, which is expected to be at mid-cell and the division site; the expected bands (rings) of fluorescence could be clearly detected for all the fusions tested, demonstrating that all the fluorescent proteins can be expressed in *H. volcanii* and used as tools for protein localization studies (Figure 2B).

**Figure 2:**
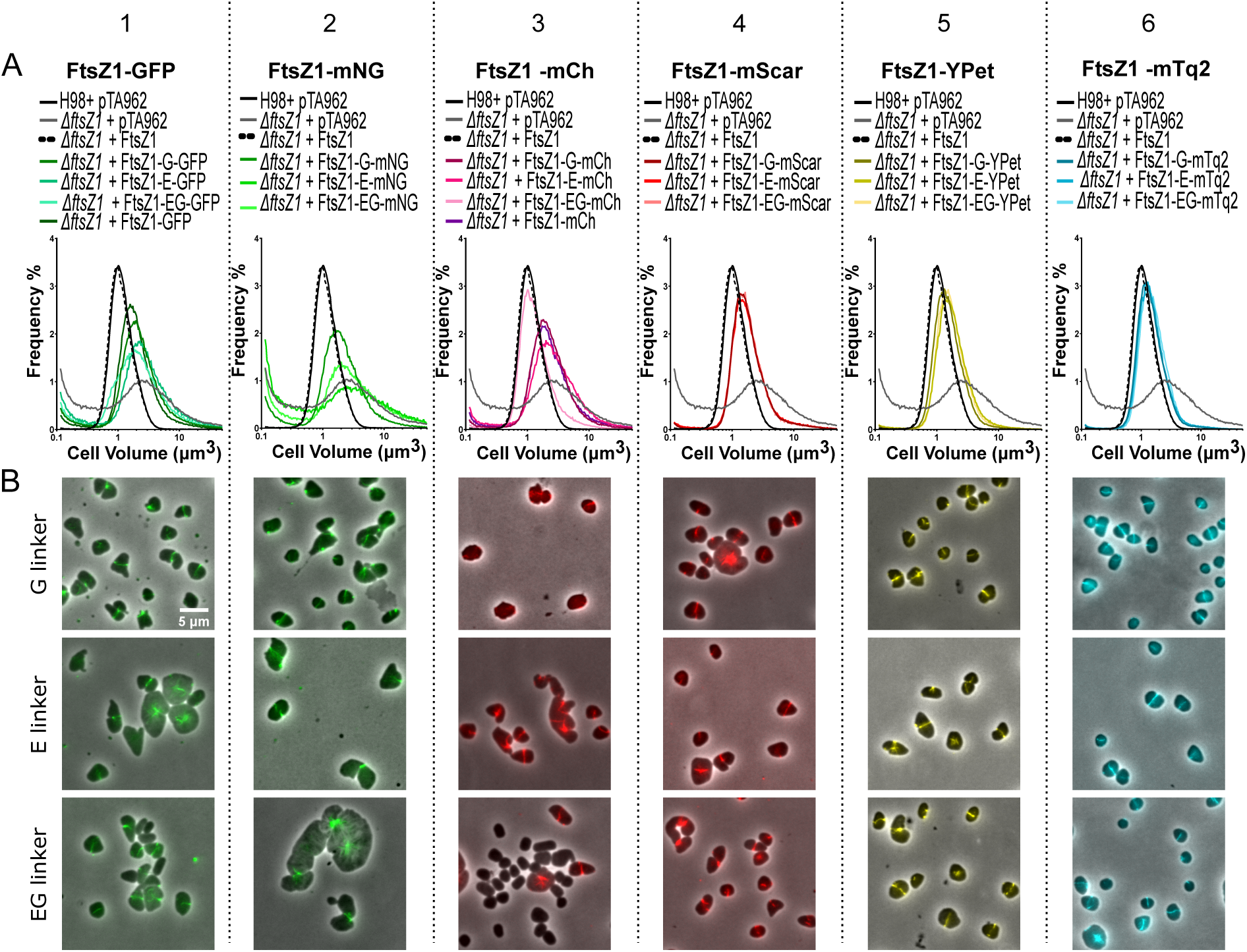
Screening of various FtsZ1-FP fusions for function in cell division and subcellular localisation. (A) Plasmids containing the indicated fusions were assessed for complementation of the cell division defect in *H. volcanii* ID76 (Δ*ftsZ1*) as assessed by Coulter cytometry cell volume distributions. (B) Composite phase-contrast and fluorescence microscopy of strains carrying the indicated FtsZ1-FP fusions. For comparison, the same datasets are shown in all graphs of the wild type (H98 + pTA962 vector only), Δ*ftsZ1* + untagged FtsZ1, and Δ*ftsZ1* control strains.

Cell volume distributions (Figure 2A) showed that none of the newly constructed FtsZ1-linker-GFP fusions nor the mNeonGreen fusions fully complemented the Δ*ftsZ1* cell volume defect; most showed only a partial rescue of the cell-size defect compared to the deletion strain control and were not improved over the original FtsZ1-GFP fusion (Figure 2A-1, 2A-2). Cells expressing GFP and mNeonGreen fusions also presented abnormal cell size and morphology by microscopy, consistent with a cell division defect, although fluorescent bands (rings) were observed at midcell or at apparent division constriction sites in most of the cells (Figure 2B-1, B-2), indicating that these GFP and mNeonGreen tags impair other activities of FtsZ1 (beyond basic midcell localization) in cell division.

The mCherry C-terminal fusions also showed partial rescue of the cell size defect of Δ*ftsZ1* (Figure 2A-3). FtsZ1-EG-mcherry appeared to largely complement the cell volume defect of *ΔftsZ1* strain, but the microscopy revealed that FtsZ1-EG-mCherry was expressed in only a small fraction of cells in this strain (Figure 2B-3) and western blots indicated the fusion was partially cleaved (Supplementary Figure 2), strongly suggesting that the apparent complementation was due to expression defects in this transformant or strain.

**Figure 3:**
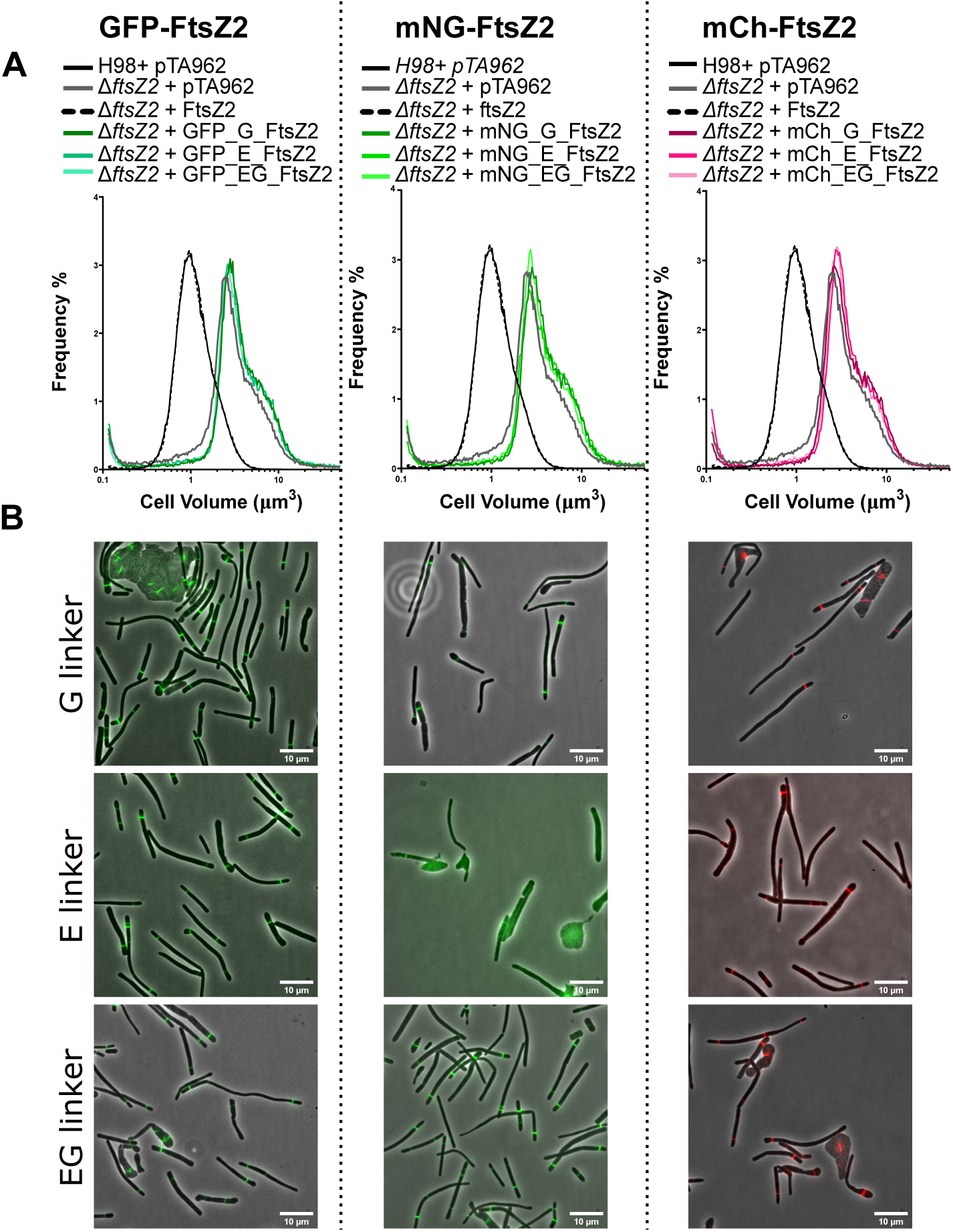
FtsZ2 N-terminus fusions localise as midcell rings but do not promote cell division. (A) Comparison of cell volume distributions measured by Coulter cytometry between WT and Δ*ftsZ2* expressing various N-terminal FP-FtsZ2 fusions. (B) Localization of various N-terminus FtsZ2 fusions in Δ*ftsZ2* strains. Overlay of Phase contrast and fluorescence channels. Scale bars = 10 µm.

All the FtsZ1 fusions and linker combinations with mScarlet-I, YPet and mTurquoise2 showed improved complementation of the cell size defect in Δ*ftsZ1* (Figure 2A4-6, 2B4-6), although not to the full degree of complementation seen with the untagged FtsZ1 plasmid control. There were no clear phenotypic differences between the three different linkers with each tag, showing that the type of FP attached to FtsZ1 had a greater impact on the function of FtsZ1 C-terminal fusions than the properties of the linker in these cases.

### Screening C- and N-terminal fluorescent protein fusions to FtsZ2 for function in H. volcanii

We next investigated FtsZ2-FP fusions, which have previously been found to dynamically localise at midcell but are non-functional in cell division [21]. FtsZ2 was cloned into the proximal position into plasmids containing the C-terminal fluorescent proteins (*i.e.*, pHVID4 to pHVID21, Figure 1B) generating 18 combinations of linker and FP fusions (plasmids pHVID58 to pHVID75) and their functionality was assessed. Despite all fusions displaying fluorescent signals and showing no or very minor proteolytic cleavage, none of the C-terminal fusion combinations complemented the Δ*ftsZ2* mutant (Supplementary Figure 3) and they showed no clear improvement compared to the previously published FtsZ2-GFP [21].

We then tested whether the fusion of the FP at the N-terminal position of FtsZ2 might improve its function. We constructed all the combinations of N-terminal fusions by cloning FtsZ2 into the distal position in the sets of plasmids containing the N-terminal fluorescent proteins (*i.e.*, pHVID22 to pHVID39, Figure 1B), thus generating 18 combinations of FtsZ2, linker and FP fusions (plasmids pHVID77 to pHVID93), and we tested their ability to complement a Δ*ftsZ2* background (Figure 3). The GFP-, mNeonGreen- and mCherry-FtsZ2 fusions showed a clear fluorescent signal (Figure 3B), but they failed to complement Δ*ftsZ2* (Figure 3A), whereas the plasmids containing mScarlet-I, YFP and mTurquoise-2 placed on the N-terminus did not show fluorescence; the same issue was observed when testing these FP N-terminal fusions with MinD4 which is known to be functional with GFP fused at its N-terminus (data not shown) [34]. On further investigation, western blotting showed that the N-terminal GFP, mNeonGreen and mCherry fusions are produced intact and at concentrations similar to or greater than the untagged FtsZ2, but, consistent with the microscopy, the mScarlet-I and mTurquoise2 fusions were barely detectable compared to expression of untagged FtsZ2, and no expression of tagged or cleaved YPet fusions was detected, consistent with the lack of fluorescence signal from these fusion proteins detected by microscopy.

### Screening of CetZ1-FP functions in H. volcanii motility and cell shape

To investigate the utilisation of the vector system in alternative functions with different expected localisation patterns, we chose the cytoskeletal protein CetZ1, which is a tubulin superfamily homologue found in archaea and is essential for rod cell development and normal swimming motility [10]. Previous work showed that overproduction of untagged CetZ1 stimulated rod formation, however, similar production of CetZ1 C-terminal fusions to smRS-GFP and mCherry (separated by a short linker of 2 amino acids, GS) did not stimulate rod-formation, suggesting the fusions were not completely functional. Nevertheless, these fusions did not interfere with rod formation in a wild-type background and showed dynamic localisation patterns consistent with the role of CetZ1 in rod cell formation. Here, we sought to identify CetZ1 fluorescent fusions with improved function, and thus demonstrate the use of the current vector system with an alternative protein, by testing the fusions for their ability to support motility and rod formation in a strain background carrying an in-frame knockout of *cetZ1*. As may be seen in Figure 4A, the constructs carrying CetZ1 fusions with E-linkers, or the mScarlet or YPet fluorescent proteins, did not significantly improve motility in the Δ*cetZ1* background, compared to the empty vector control. For the GFP, mCherry, and mTurquoise2 fusions, with either the EG- or G-linkers, we observed a substantial restoration of motility, whereas for mNeonGreen, only the combination with the G-linker somewhat restored motility. Measurement of motility halo diameter with replicate experiments (Figure 4B) showed that the strain carrying CetZ1-EG-mTq2 retained almost complete function in motility (∼97%) compared to the that with the untagged *cetZ1*.

**Figure 4:**
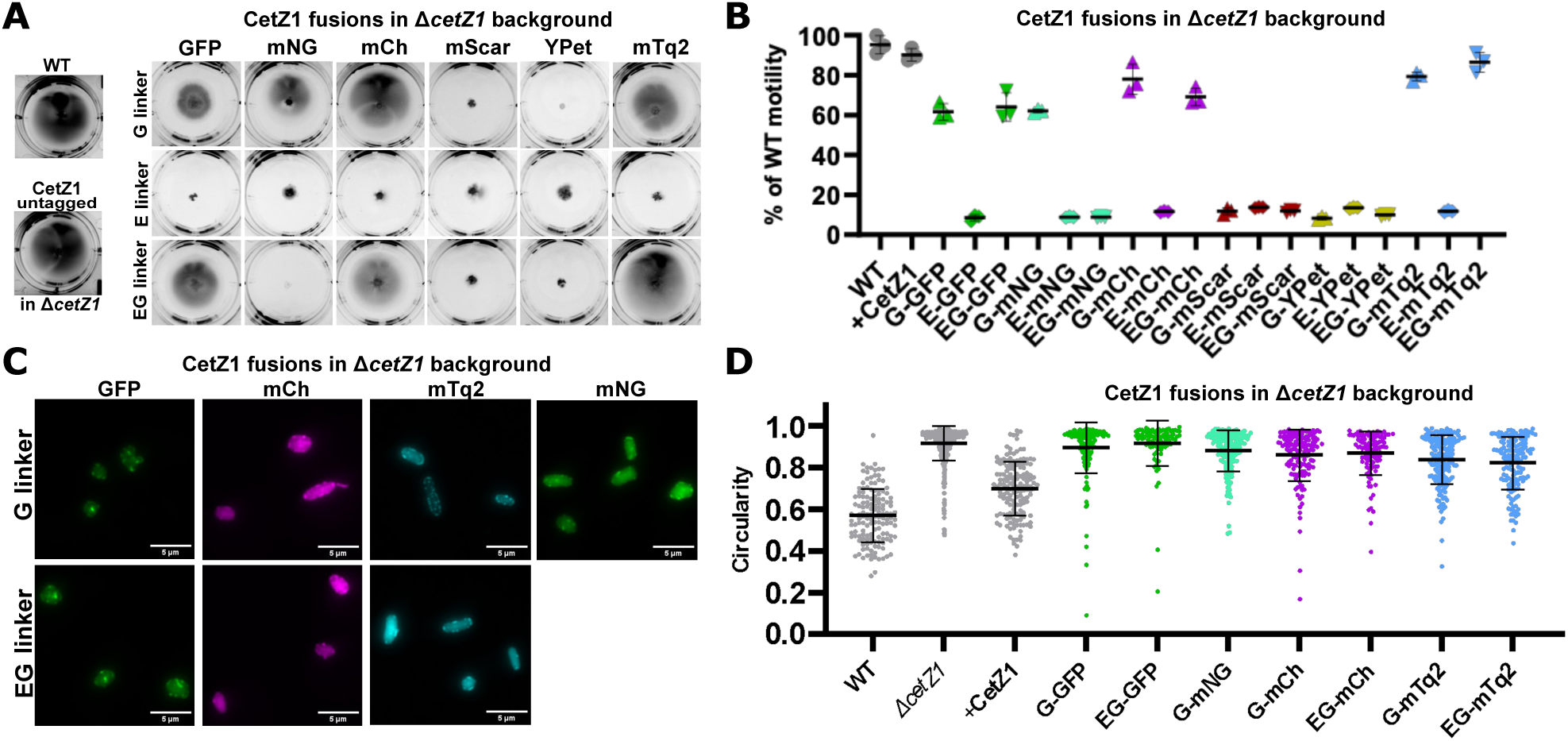
Screening Δ*cetZ1* complementation by CetZ1-FP. CetZ1 C-terminal fusions and untagged CetZ1 control were produced in the Δ*cetZ1* genetic background (ID181). (A) Representative images of swimming motility assays in Hv-Cab medium (0.3 % agar) supplemented with 2 mM Trp inducer to maintain production of the indicated proteins. A wild-type reference strain was included (H26 + pTA962). (B) Quantification of motility halo diameter from three replicate halos (mean and SEM overlaid). (C) Representative fluorescence microscopy images of CetZ1 fusions expressed in Δ*cetZ1* (Hv-Cab + 2 mM Trp) which at least partially complemented the Δ*cetZ1* motility defect. Similar patterns were observed for proteins that failed to complement Δ*cetZ1* (not shown). (D) Quantification of cell circularity of Δ*cetZ1* expressing selected CetZ1 fusion proteins. Individual points represent individual cells, and data shown is representative of two replicates. Bars represent mean ± standard deviation.

For the CetZ1 fusions that at least partially restored motility, we next observed their subcellular localisation and measured their capacity to complement the rod-shape defect in the Δ*cetZ1* background. As may be seen in Figure 4C, all these strains showed fluorescent signal within the cells and similar localization patterns compared to the previously reported GFP and mCherry fusions [10]. Western blotting detection of CetZ1 (Figure S5) showed generally elevated productions levels compared to the untagged protein, and in some cases minor additional bands were observed, consistent with modified forms appearing such as after potential proteolytic cleavage or alternative translation termination events. Several of these fusion proteins, notably the relatively stable G- and EG-linked mCherry and mTurquoise2 fusions, provided substantial rescue of the rod-development defect of the *ΔcetZ1* background as detected via a reduced mean cell circularity in the population (Figure 4D). The GFP and mNeonGreen constructs analysed showed partial complementation (Figure 4C and 4D) like the original CetZ1-GFP (not shown) [10]. These results corresponded very well to motility (compare to Figure 4B) and indicate that CetZ1-EG-mTq2 was overall the most functional fusion protein. They also demonstrate the relevance of screening a range of different combinations of linkers and fluorescent proteins when searching for fusion proteins with optimal function in *H. volcanii*.

### Testing of the dual expression module of the vector set with co-expression of FtsZ1 and FtsZ2

As shown in Figure 1C, the vector design facilitates construction of plasmids for expression of two tandemly positioned genes or fusion genes under the regulation of one promoter, p.*tnaA*. To evaluate the function of the tandem expression system, we constructed two plasmids expressing tagged FtsZ1 and FtsZ2 in two alternate configurations: pHVID127, expressing *ftsZ1*-EG-*mTurquoise-2*::*mCherry*-EG-*ftsZ2* and pHVID128, expressing *mCherry*-EG-*ftsZ2::ftsZ1*-EG-*mTurquoise-2*. Fluorescence microscopy of strains carrying these plasmids in a wild-type *H. volcanii* background (Figure 5A) showed mid-cell ring localization in both fluorescence channels in the two strains, consistent with the roles of the proteins in division (Figure 5A). To determine whether the position of the genes in the synthetic operon could affect their expression, fluorescence intensity was measured as the mean pixel intensity per cell (Figure 5B, 5C), and the ratio between mCherry and mTurquoise2 total pixel intensities in individual cells was then determined for each strain (Figure 5D). This showed that mCherry-EG-FtsZ2 was expressed at similar levels, whether it was in the first or second position in the synthetic operon (Figure 5B). However, FtsZ1-EG-mTurquoise2 expression was significantly higher from pHVID128, where its ORF is in the first position (Figure 5A, 5C). We conclude that the relative positioning of tandem genes encoding fluorescent proteins influences their expression, depending on the gene.

**Figure 5:**
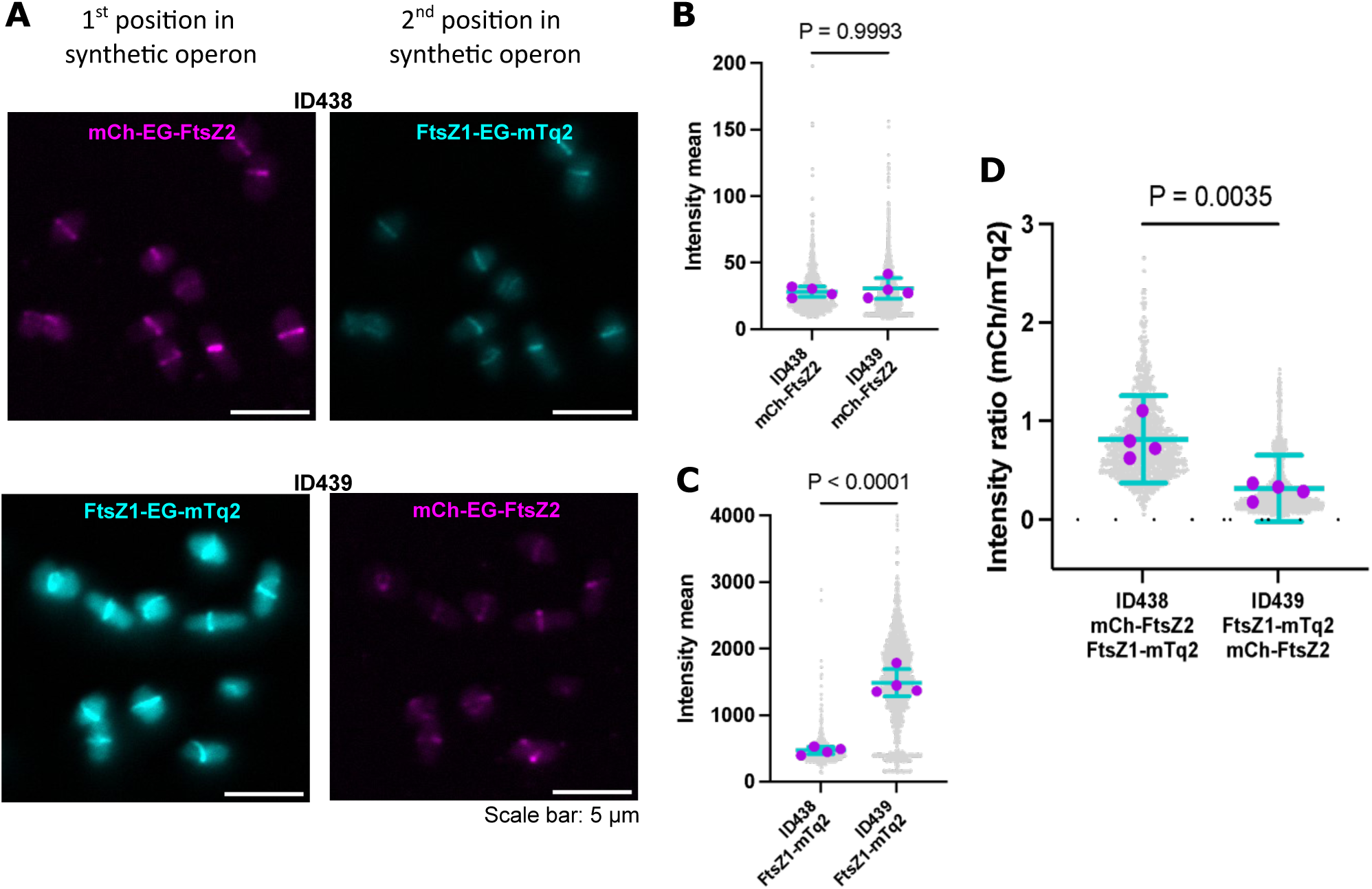
Tandem expression of FtsZ1-EG-mTq2 and mCh-EG-FtsZ2. (A) *H. volcanii* strains ID438 (expressing mCh-EG-FtsZ2 from the first gene position and FtsZ1-EG-mTq2 from the second gene position) and ID439 (expressing the genes in the reverse order) sampled during steady mid-log growth in Hv-Cab + 0.2 mM Trp, and imaged using fluorescence microscopy using identical imaging conditions and processing. Representative images of FtsZ1 and FtsZ2 in ID438 and ID439. The mean fluorescence intensities in individual cells for the mCherry (B) and mTurquoise2 (C) channels were measured for each individual cell in both strains. (D) Then, the ratio between mCherry total intensity and mTurquoise2 intensity was calculated for each individual cell and plotted for ID438 and ID439. For (B), (C), and (D), grey points represent all measured individual cells from four biological replicates, magenta points indicate the average of each biological replicate, and cyan error bars represent the mean and standard deviation of biological replicate averages. P-values from t-tests are shown above each graph.

## Discussion

Cell biology and spatiotemporal cell organisation studies in archaea have gained interest in recent years, taking advantage of genetically encoded FP fusions for dynamic subcellular localization studies, which have contributed significantly to the identification of new archaeal functions and mechanisms [10, 19, 20, 34]. However, as seen with protein fusions in bacterial or eukaryotic systems [14, 35], finding functional FP fusions in archaea can be challenging and time-consuming [10, 19, 21]. We have devised *H. volcanii* plasmid vectors based on the commonly used pHV2 replication origin and tryptophan-inducible promoter [5, 36] that facilitate construction and functional testing and screening of proteins and fusion proteins, in both single and dual (tandem) configurations, for *in vivo* studies of protein function and localization.

We demonstrated uses of this system using three *H. volcanii* target proteins and, for designing fusion proteins, we showed the importance of screening a range of combinations of linkers and fluorescent proteins in identifying functional fusion proteins in *H. volcanii*. Biological functionality could not be accurately predicted in advance, consistent with observations and recommendations for bacterial and eukaryotic systems [14, 35]. It is remarkable that the various fluorescent proteins, which are generally highly homologous, showed substantial variation in their functional consequences on the tagged protein, further highlighting the importance of functional testing of each new fusion protein variant.

Despite these varying effects on function, the subcellular localization patterns of each target protein with various linker/tag combinations appeared much more consistent; the linkers/tags are expected to affect any of the multiple protein activities other than subcellular localization that contribute to their function. An apparent exception to this included FtsZ1-mNeonGreen fusions, which failed to rescue the cell division defect in the Δ*ftsZ1* background and their localization also appeared anomalous. However, in this case, at least some of the aberrant localization is likely to be a consequence of the substantial enlargement and deformation of these cells, rather than just an intrinsic inability of the protein to assemble at midcell. This is consistent with previous observations of FtsZ1-mCherry in other cell division mutant backgrounds [21].

Although we were able to identify highly functional FtsZ1 and CetZ1 fusion proteins, functional screening of the full sets of FtsZ2 C- and N-terminal fusions identified no improvement of the cell division defects of a Δ*ftsZ2* mutant, even though most of these proteins showed the capacity to localize as distinct rings in the enlarged filamentous cells. It is possible that an alternative approach involving screening of internally positioned tags would identify improved FtsZ2-FP functionality, as seen with *E. coli* FtsZ, for example [17, 37].

During our investigations of various fusion proteins and dual (tandem) expression plasmids that facilitate multicolour imaging in *H. volcanii*, we observed several gene position dependent effects on expression. Unexpectedly, three of the codon-optimized FPs we generated here (mScarlet-I, YPet, and mTurquoise2) were expressed at extremely low or undetectable levels (by western blots and fluorescence microscopy) only when they were positioned in the N-terminal position with respect to the tagged protein; whereas other N-terminal FP fusions to the same *H. volcanii* proteins were expressed well and localize specifically. Other observations have indicated that lack of expression in these constructs also occurs when the fusion is in the second position in a tandem construct. These findings suggest that gene expression in *H. volcanii* can be strongly dependent on the sequence of the N-terminal coding region, and we speculate this could relate to mRNA folding or stability that has a significant impact on amount of protein translated. Furthermore, we observed differences in the relative level of expression of certain fusions depending on whether they are in the first or second position of tandem constructs (Figure 5). Thus, the presence or structure of untranslated regions in these cases also appears to contribute significantly to levels of the final protein products. Further studies of how the predicted structure and stability of mRNA around the start codon influence gene expression in *H. volcanii* should facilitate further improvements to protein production control and for high-level overproduction in research or biotechnology applications of archaea.

## Acknowledgements

The authors acknowledge technical and facilities management support from Louise Cole and Amy Bottomley, and use of equipment in the Microbial Imaging Facility, Faculty of Science, University of Technology Sydney. This project was supported by the Australian Research Council (FT160100010 and DP160101076).

