## Supplementary Information for "A vector system for single and tandem expression of cloned genes and multi-colour fluorescent tagging in *Haloferax volcanii*"

**Supplementary Figures**

1. Construction of pIDJL114_noXbaI and p.*tnaA**
2. FtsZ1-FP fusion protein western blots
3. FtsZ2-FP (C-terminal) fusions and western blots
4. FP-FtsZ2 (N-terminal) fusions and western blots
5. Dual localization/western blots
6. Western for CetZ1-FP fusions

**Supplementary Tables**

1. Plasmids and oligonucleotides
2. Cloned Gene Sequences

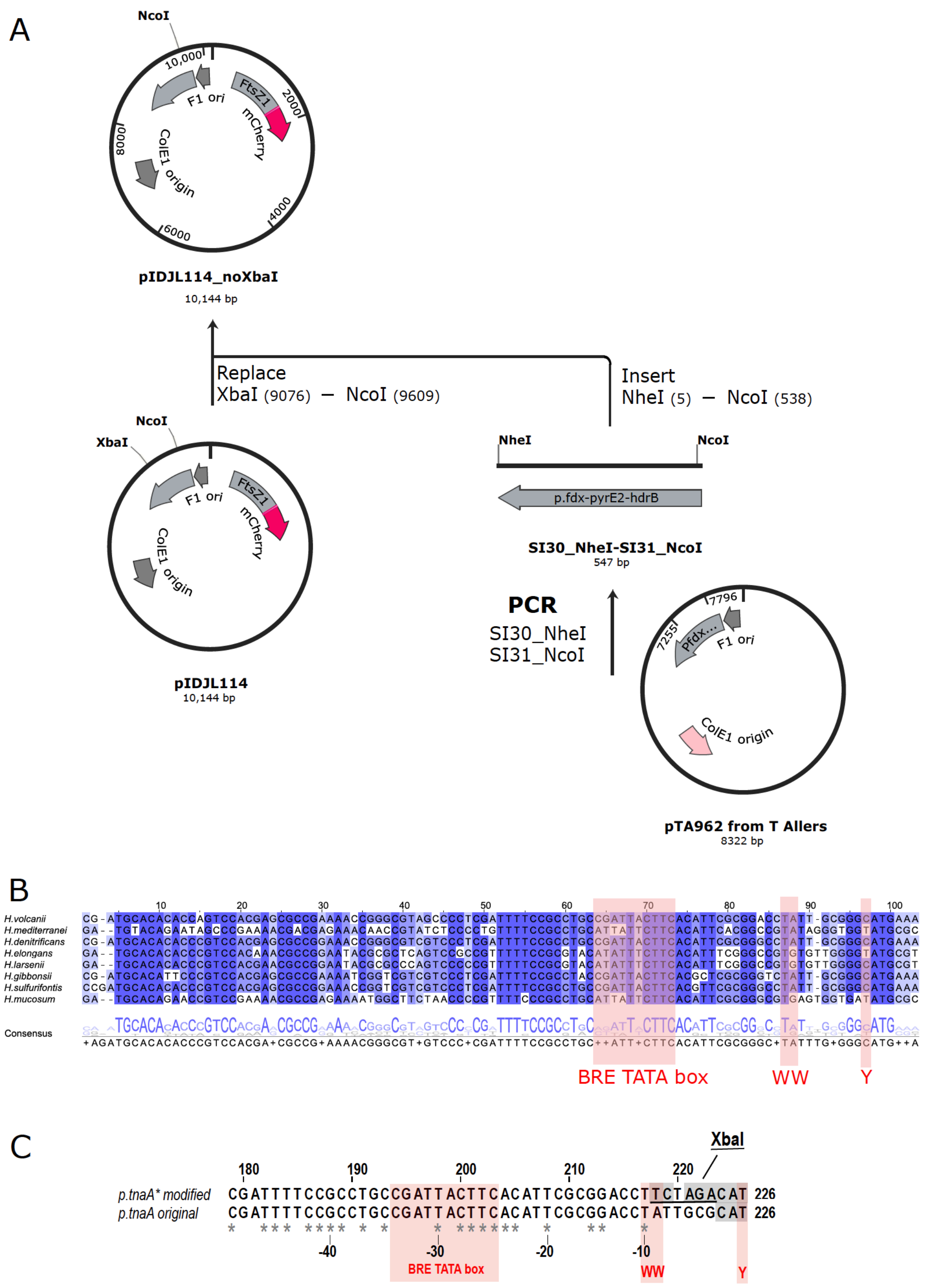


**Supplementary Figure 1. Plasmid preparation steps and modification of the p.*tnaA* promoter to accommodate an XbaI site to facilitate tandem cloning.** (A) We began with pIDJL114 [1], a pTA962-derived vector containing a FtsZ1-mCherry fusion cloned between the BamHI and NotI sites of pTA962 thereby avoiding one XbaI site in the original MCS of pTA962. To remove the remaining XbaI site located between the *hdrB* and *pyrE2* genes, we used a PCR amplified the *pyrE2* fragment from pTA962 with primers SI30 and SI31, with NheI/NcoI ends. This was used to replace the equivalent XbaI/NcoI fragment within pIDJL114, generating a plasmid containing no XbaI sites (pIDJL114 NoXbaI), which conserves the spacing between the tandem *hdrB* and *pyrE2* ORFs in pTA962. (B) A multiple alignment of selected Haloferacales p.*tnaA* promoter regions. ClustalW alignments of 0.1 kb regions upstream of *tna*A open reading frames (the first two codons are shown) from the indicated Haloferacales species were visualized with Jalview. Pink shading represents the predicted regions containing basal transcription regulatory elements, including the BRE, TATA box, WW and pyrimidine (Y) -1 motifs [2]. These alignments were used to assess whether the introduced XbaI site (at -9 to -4) in the p.*tnaA* promoter of *H. volcanii* might affect transcription initiation; the lower conservation in the region -9 to -4 between the WW and Y motifs suggested that modifications in this region would not significantly disturb transcription. (C) DNA sequence alignment of the original [3] and modified p.*tnaA* promoters. Locations of the expected promoter consensus elements (BRE, TATA box, WW, and Y motifs) are shown by pink shading. The XbaI site introduced in this study is underlined. Grey shading indicates nucleotides differing from the wild-type chromosomal p.*tnaA* promoter. Asterisks indicate nucleotides conserved from (B).

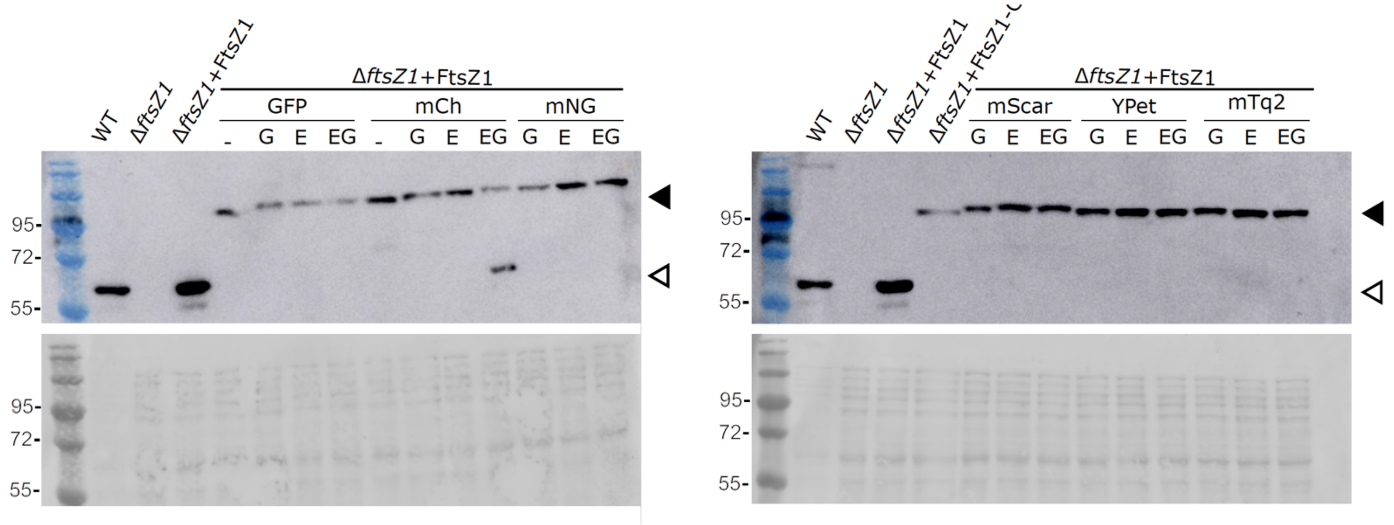


**Supplementary Figure 2.** Western blotting of whole-cell protein lysates to detect production of FtsZ1 in strains containing the indicated FtsZ1-linker-FP combinations. Approximate marker sizes are indicated (kDa). Controls include H98 (WT), ID76 (Δ*ftsZ1*), and ID76 with native, untagged FtsZ1 expressed from a plasmid. Upper panels represent anti-FtsZ1 detection, and the lower panels show Ponceau S total protein stain of the same membranes. Triangles indicate the approximate position of native FtsZ (lower band) and FtsZ1-FP (upper band).

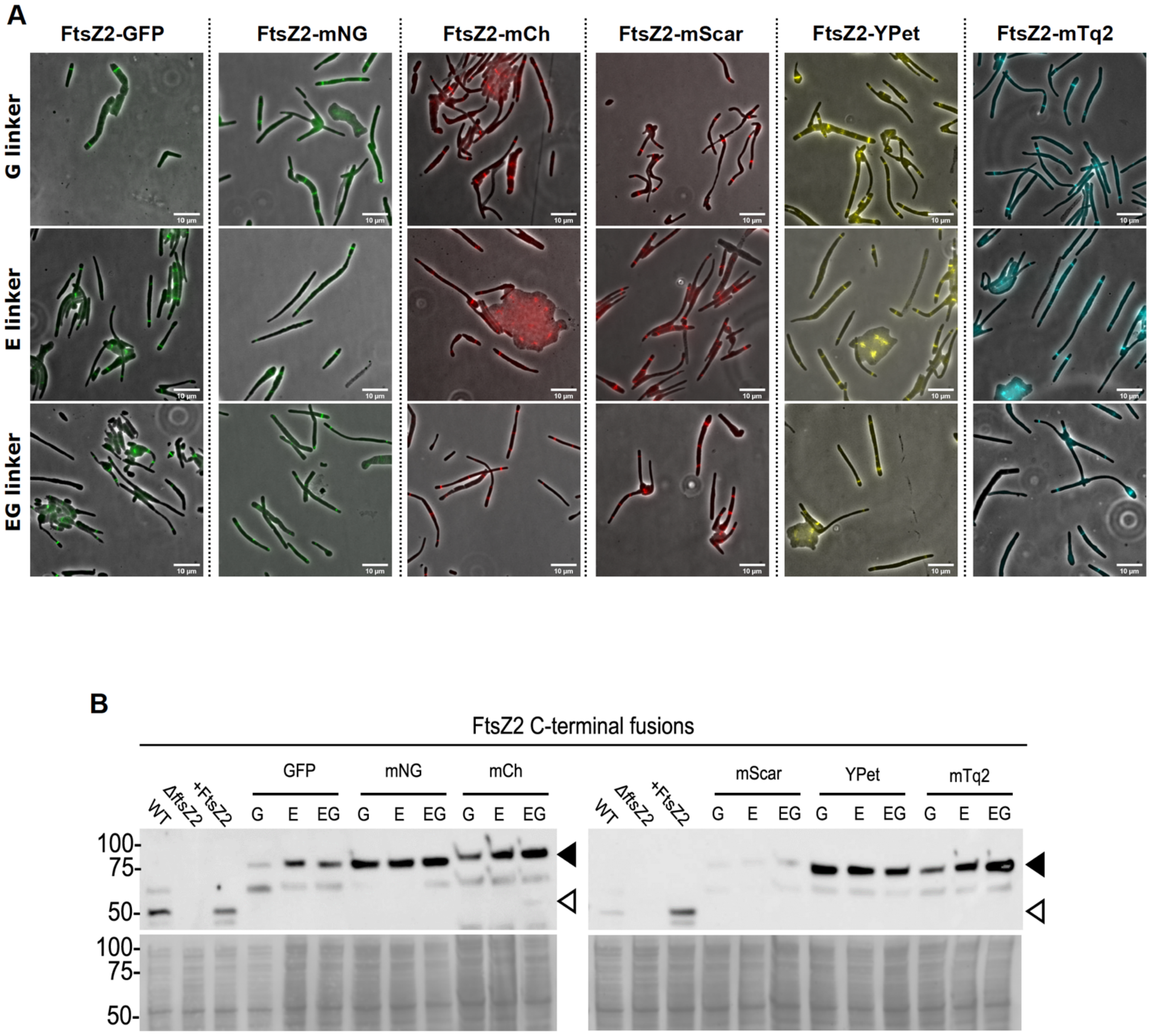


**Supplementary Figure 3**: FtsZ2-FP C-terminal fusions can localize or form rings but do not promote cell division. (A) Localization of various FtsZ2 C-terminus fusions in Δ*ftsZ2* strains. Composite images of phase-contrast and fluorescence with the indicated combinations of fluorescent proteins and linker. Scale bars = 10 µm. (B) Western blot detection of FtsZ2 in whole cell lysates of the indicated *H. volcanii* strains (ID77 (Δ*ftsZ2*) background, except for WT). Upper – Anti-FtsZ2 antibody. Lower – Ponceau S total protein stain. Marker sizes (kDa) are indicated on left of images. Approximate position of the fusion proteins and untagged FtsZ2 are indicated by filled and empty triangles, respectively.

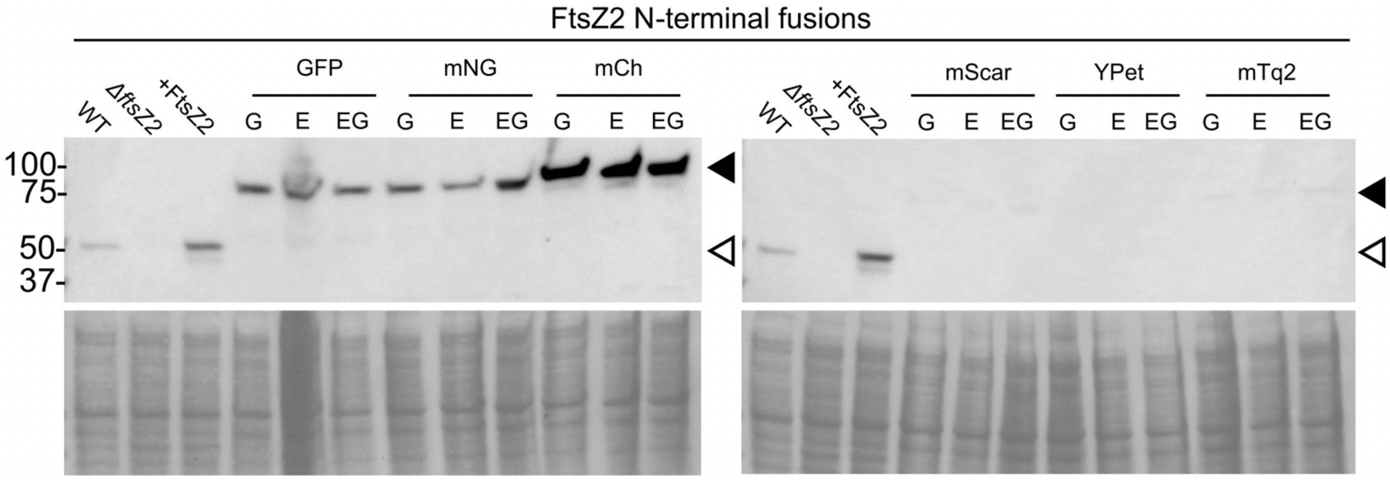


**Supplementary Figure 4.** Western blot detection of FtsZ2 in whole cell lysates of the indicated *H. volcanii* strains (ID77 (Δ*ftsZ2*) background, except for WT) carrying the indicated FPs fused to the N-terminus of FtsZ2. Upper panels, anti-FtsZ2 antibody. Lower panels, Ponceau S total protein stain. Marker sizes (kDa) indicated on left of images.)

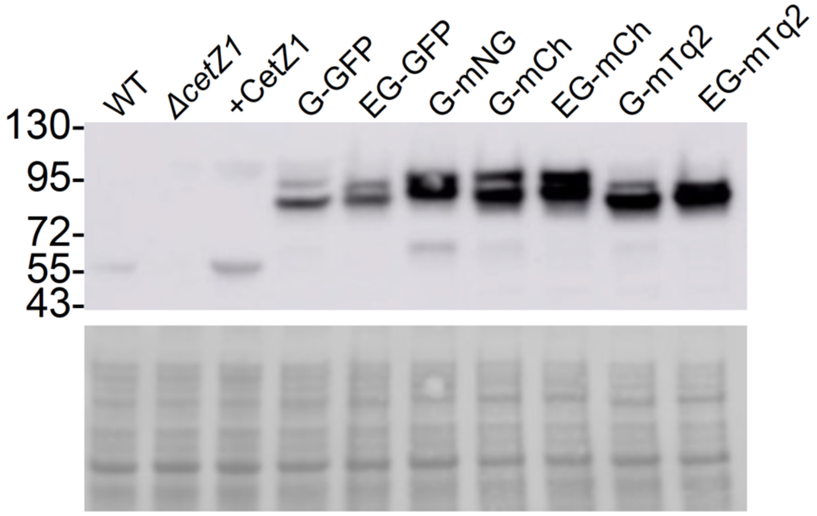


**Supplementary Figure 5.** Western blotting detection of CetZ1 in Δ*cetZ1* strains expressing the indicated CetZ1-FP fusions. Upper panel – CetZ1 probed membrane. Lower panel – Ponceau S total protein-stained membrane. The major CetZ1-probed band appearing in the lanes representing fusion proteins is expected to represent the relative amount of full-length fusion protein in vivo. Replicate western blots (not shown) showed some variation in the degree of additional minor banding in some CetZ1-FP strains, suggesting that the fusions might be sensitive to modification in different culture samples or during protein extraction.

**Supplementary Table 1.** Plasmids and oligonucleotides used for their construction.

| Plasmid | Description/function | Sequences of synthetic DNA used for construction (5' to 3'), where relevant | Source |
|---|---|---|---|
| pTA962 | *p.tnaA* expression vector for *H. volcanii. E. coli shuttle plasmid.* | | [4] |
| pIDJL40 | pTA962 with smRS*gfp* (BamHI-NotI) | | [5] |
| pTA962-*ftsZ1* | *P.tna* control of *ftsZ1* | | [1] |
| pTA962-*ftsZ2* | *P.tna* control of *ftsZ2* | | [1] |
| pIDJL40-*ftsZ1* | *P.tna* control of *ftsZ1-gfp* | | [1] |
| pIDJL40-*ftsZ2* | *P.tna* control of *ftsZ2-gfp* | | [1] |
| pIDJL114 | *P.tna* control of *ftsZ1-mCherry* | | [1] |
| pIDJL115 | *P.tna* control of *ftsZ2-mCherry* | | [1] |
| pIDJL114_NoXbaI | *Contains silent mutation to remove an XbaI restriction site between pyrE2 and hdrB* | *pyrE2//CTGGCCGACGCCGACGGCTAAGCTAGATGAGCGGCGAGGAGCTTTCTG//hdrB* | This study |
| pHVID1 | P.tnaA*_Elinker | Synthetised fragment cloned into pIDJL114_XbaIremoved digested ApaI/NotI | This study |
| pHVID2 | P.tnaA*_Glinker | Synthetised fragment cloned into pIDJL114_XbaIremoved digested ApaI/NotI | This study |
| pHVID3 | P.tnaA*_EGlinker | Synthetised fragment cloned into pIDJL114_XbaIremoved digested ApaI/NotI | This study |
| pHVID4 | P.tnaA*_G_GFP | Synthetized fragment cloned into pHVID2 EcoRI/NheI | This study |
| pHVID5 | P.tnaA*_G_$^{Hv}$mNG | Synthetized fragment cloned into pHVID2 EcoRI/NheI | This study |
| pHVID6 | P.tnaA*_G_$^{Hv}$mCh | Synthetized fragment cloned into pHVID2 EcoRI/NheI | This study |
| pHVID7 | P.tnaA*_G_$^{Hv}$mScar-I | Synthetized fragment cloned into pHVID2 EcoRI/NheI | This study |
| pHVID8 | P.tnaA*_G_$^{Hv}$YPet | Synthetized fragment cloned into pHVID2 EcoRI/NheI | This study |
| pHVID9 | P.tnaA*_G_$^{Hv}$mTq-2 | Synthetized fragment cloned into pHVID2 EcoRI/NheI | This study |
| pHVID10 | P.tnaA*_E_GFP | Synthetized fragment cloned into pHVID1 EcoRI/NheI | This study |
| pHVID11 | P.tnaA*_E_$^{Hv}$mNG | Synthetized fragment cloned into pHVID1 EcoRI/NheI | This study |
| pHVID12 | P.tnaA*_E_$^{Hv}$mCh | Synthetized fragment cloned into pHVID1 EcoRI/NheI | This study |
| pHVID13 | P.tnaA*_E_$^{Hv}$mScar-I | Synthetized fragment cloned into pHVID1 EcoRI/NheI | This study |
| pHVID14 | P.tnaA*_E_$^{Hv}$YPet | Synthetized fragment cloned into pHVID1 EcoRI/NheI | This study |
| pHVID15 | P.tnaA*_E_$^{Hv}$mTq-2 | Synthetized fragment cloned into pHVID1 EcoRI/NheI | This study |
| pHVID16 | P.tnaA*_EG_GFP | Synthetized fragment cloned into pHVID3 EcoRI/NheI | This study |
| pHVID17 | P.tnaA*_EG_$^{Hv}$mNG | Synthetized fragment cloned into pHVID3 EcoRI/NheI | This study |
| pHVID18 | P.tnaA*_EG_$^{Hv}$mCh | Synthetized fragment cloned into pHVID3 EcoRI/NheI | This study |
| pHVID19 | P.tnaA*_EG_$^{Hv}$mScar-I | Synthetized fragment cloned into pHVID3 EcoRI/NheI | This study |
| pHVID20 | P.tnaA*_EG_$^{Hv}$YPet | Synthetized fragment cloned into pHVID3 EcoRI/NheI | This study |
| pHVID21 | P.tnaA*_EG_$^{Hv}$mTq-2 | Synthetized fragment cloned into pHVID3 EcoRI/NheI | This study |
| pHVID22 | P.tnaA*_GFP_G | PCR cloned into pHVID2 NdeI/BamHI<br>SI34_NdeI GTAT**CATATG**AGTAAAGGAGAAGAACTTTT<br>SI35_BamHI GTAC**GGATCC**TTTGTATAGTTCATCCATGC | This study |
| pHVID23 | P.tnaA*_$^{Hv}$mNG_G | PCR cloned into pHVID2 NdeI/BamHI<br>SI38_NdeI TGTG**CATATG**GTCTCGAAGGGCGAGGAAGA<br>SI118_BamHI GTAT**GGATCC**CTTGTAGAGCTCGTCCATGCC | This study |
| pHVID24 | P.tnaA*_$^{Hv}$mCh_G | PCR cloned into pHVID2 NdeI/BamHI<br>SI36_NdeI TGAG**CATATG**GTCTCGAAGGGCGAGGAGGA<br>SI37_BamHI TGAT**GGATCC**CTTGTAGAGCTCGTCCATGC | This study |
| pHVID25 | P.tnaA*_$^{Hv}$mScar-I_G | PCR cloned into pHVID2 NdeI/BamHI<br>SI40_NdeI GTAG**CATATG**GTCTCGAAGGGCGAGGCCGT<br>SI41_BamHI GTAT**GGATCC**CTTGTAGAGCTCGTCCATGC | This study |
| pHVID26 | P.tnaA*_$^{Hv}$YPet_G | PCR cloned into pHVID2 NdeI/BamHI<br>SI44_NdeI GTAT**CATATG**TCGAAGGGCGAGGAGCTCTT<br>SI45_BamHI GTTA**GGATCC**CTTGTAGAGCTCGTTCATGC | This study |
| pHVID27 | P.tnaA*_$^{Hv}$mTq-2_G | PCR cloned into pHVID2 NdeI/BamHI<br>SI42_NdeI GTAT**CATATG**GTCTCGAAGGGCGAGGAGCT<br>SI43_BamHI GTAT**GGATCC**CTTGTAGAGCTCGTCCATG | This study |
| pHVID28 | P.tnaA*_GFP-E | PCR cloned into pHVID1 NdeI/BamHI<br>SI34_NdeI GTAT**CATATG**AGTAAAGGAGAAGAACTTTT<br>SI35_BamHI GTAC**GGATCC**TTTGTATAGTTCATCCATGC | This study |
| pHVID29 | P.tnaA*_$^{Hv}$mNG_E | PCR cloned into pHVID1 NdeI/BamHI<br>SI38_NdeI TGTG**CATATG**GTCTCGAAGGGCGAGGAAGA<br>SI118_BamHI GTAT**GGATCC**CTTGTAGAGCTCGTCCATGCC | This study |
| pHVID30 | P.tnaA*_$^{Hv}$mCh_E | PCR cloned into pHVID1 NdeI/BamHI<br>SI36_NdeI TGAG**CATATG**GTCTCGAAGGGCGAGGAGGA<br>SI37_BamHI TGAT**GGATCC**CTTGTAGAGCTCGTCCATGC | This study |
| pHVID31 | P.tnaA*_$^{Hv}$mScar-I_E | PCR cloned into pHVID1 NdeI/BamHI<br>SI40_NdeI GTAG**CATATG**GTCTCGAAGGGCGAGGCCGT<br>SI41_BamHI GTAT**GGATCC**CTTGTAGAGCTCGTCCATGC | This study |
| pHVID32 | P.tnaA*_$^{Hv}$YPet_E | PCR cloned into pHVID1 NdeI/BamHI<br>SI44_NdeI GTAT**CATATG**TCGAAGGGCGAGGAGCTCTT<br>SI45_BamHI GTTA**GGATCC**CTTGTAGAGCTCGTTCATGC | This study |

| | | | |
|---|---|---|---|
| pHVID33 | P.tnaA*_$^{Hv}$mTq-2_E | PCR cloned into pHVID1 NdeI/BamHI<br>SI42_NdeI GTAT**CATATG**GTCTCGAAGGGCGAGGAGCT<br>SI43_BamHI GTAT**GGATCC**CTTGTAGAGCTCGTCCATG | This study |
| pHVID34 | P.tnaA*_GFP_EG | PCR cloned into pHVID3 NdeI/BamHI<br>SI34_NdeI GTAT**CATATG**AGTAAAGGAGAAGAACTTTT<br>SI35_BamHI GTAC**GGATCC**TTTGTATAGTTCATCCATGC | This study |
| pHVID35 | P.tnaA*_$^{Hv}$mNG_EG | PCR cloned into pHVID3 NdeI/BamHI<br>SI38_NdeI TGTG**CATATG**GTCTCGAAGGGCGAGGAAGA<br>SI118_BamHI GTAT**GGATCC**CTTGTAGAGCTCGTCCATGCC | This study |
| pHVID36 | P.tnaA*_$^{Hv}$mCh_EG | PCR cloned into pHVID3 NdeI/BamHI<br>SI36_NdeI TGAG**CATATG**GTCTCGAAGGGCGAGGAGGA<br>SI37_BamHI TGAT**GGATCC**CTTGTAGAGCTCGTCCATGC | This study |
| pHVID37 | P.tnaA*_$^{Hv}$mScar-II_EG | PCR cloned into pHVID3 NdeI/BamHI<br>SI40_NdeI GTAG**CATATG**GTCTCGAAGGGCGAGGCCGT<br>SI41_BamHI GTAT**GGATCC**CTTGTAGAGCTCGTCCATGC | This study |
| pHVID38 | P.tnaA*_$^{Hv}$YPet_EG | PCR cloned into pHVID3 NdeI/BamHI<br>SI44_NdeI GTAT**CATATG**TCGAAGGGCGAGGAGCTCTT<br>SI45_BamHI GTTA**GGATCC**CTTGTAGAGCTCGTTCATGC | This study |
| pHVID39 | P.tnaA*_$^{Hv}$mTq-2_EG | PCR cloned into pHVID3 NdeI/BamHI<br>SI42_NdeI GTAT**CATATG**GTCTCGAAGGGCGAGGAGCT<br>SI43_BamHI GTAT**GGATCC**CTTGTAGAGCTCGTCCATG | This study |
| pHVID40 | P.tnaA*_FtsZ1_G_GFP | NdeI-*ftsZ1*-BamHI from pIDJL40-*ftsZ1* cloned into pHVID4 | This study |
| pHVID41 | P.tnaA*_FtsZ1_G_$^{Hv}$mNG | NdeI-*ftsZ1*-BamHI from pIDJL40-*ftsZ1* cloned into pHVID5 | This study |
| pHVID42 | P.tnaA*_FtsZ1_G_$^{Hv}$mCh | NdeI-*ftsZ1*-BamHI from pIDJL40-*ftsZ1* cloned into pHVID6 | This study |
| pHVID43 | P.tnaA*_FtsZ1_G_$^{Hv}$mScar-I | NdeI-*ftsZ1*-BamHI from pIDJL40-*ftsZ1* cloned into pHVID7 | This study |
| pHVID44 | P.tnaA*_FtsZ1_G_$^{Hv}$YPet | NdeI-*ftsZ1*-BamHI from pIDJL40-*ftsZ1* cloned into pHVID8 | This study |
| pHVID45 | P.tnaA*_FtsZ1_G_$^{Hv}$mTq-2 | NdeI-*ftsZ1*-BamHI from pIDJL40-*ftsZ1* cloned into pHVID9 | This study |
| pHVID46 | P.tnaA*_FtsZ1_E_GFP | NdeI-*ftsZ1*-BamHI from pIDJL40-*ftsZ1* cloned into pHVID10 | This study |
| pHVID47 | P.tnaA*_FtsZ1_E_$^{Hv}$mNG | NdeI-*ftsZ1*-BamHI from pIDJL40-*ftsZ1* cloned into pHVID11 | This study |
| pHVID48 | P.tnaA*_FtsZ1_E_$^{Hv}$mCh | NdeI-*ftsZ1*-BamHI from pIDJL40-*ftsZ1* cloned into pHVID12 | This study |
| pHVID49 | P.tnaA*_FtsZ1_E_$^{Hv}$mScar-I | NdeI-*ftsZ1*-BamHI from pIDJL40-*ftsZ1* cloned into pHVID13 | This study |
| pHVID50 | P.tnaA*_FtsZ1_E_$^{Hv}$YPet | NdeI-*ftsZ1*-BamHI from pIDJL40-*ftsZ1* cloned into pHVID14 | This study |
| pHVID51 | P.tnaA*_FtsZ1_E_$^{Hv}$mTq-2 | NdeI-*ftsZ1*-BamHI from pIDJL40-*ftsZ1* cloned into pHVID15 | This study |
| pHVID52 | P.tnaA*_FtsZ1_EG_GFP | NdeI-*ftsZ1*-BamHI from pIDJL40-*ftsZ1* cloned into pHVID16 | This study |
| pHVID53 | P.tnaA*_FtsZ1_EG_$^{Hv}$mNG | NdeI-*ftsZ1*-BamHI from pIDJL40-*ftsZ1* cloned into pHVID17 | This study |
| pHVID54 | P.tnaA*_FtsZ1_EG_$^{Hv}$mCh | NdeI-*ftsZ1*-BamHI from pIDJL40-*ftsZ1* cloned into pHVID18 | This study |
| pHVID55 | P.tnaA*_FtsZ1_EG_$^{Hv}$mScar-I | NdeI-*ftsZ1*-BamHI from pIDJL40-*ftsZ1* cloned into pHVID19 | This study |
| pHVID56 | P.tnaA*_FtsZ1_EG_$^{Hv}$YPet | NdeI-*ftsZ1*-BamHI from pIDJL40-*ftsZ1* cloned into pHVID20 | This study |
| pHVID57 | P.tnaA*_FtsZ1_EG_$^{Hv}$mTq-2 | NdeI-*ftsZ1*-BamHI from pIDJL40-*ftsZ1* cloned into pHVID21 | This study |
| pHVID58 | P.tnaA*_FtsZ2_Glink_GFP | NdeI-*ftsZ2*-BamHI from pIDJL40-*ftsZ2* cloned into pHVID4 | This study |
| pHVID59 | P.tnaA*_FtsZ2_Glink_$^{Hv}$mNG | NdeI-*ftsZ2*-BamHI from pIDJL40-*ftsZ2* cloned into pHVID5 | This study |
| pHVID60 | P.tnaA*_FtsZ2_Glink_$^{Hv}$mCh | NdeI-*ftsZ2*-BamHI from pIDJL40-ftsZ2 cloned into pHVID6 | This study |
| pHVID61 | P.tnaA*_FtsZ2_Glink_$^{Hv}$mScar-I | NdeI-*ftsZ2*-BamHI from pIDJL40-ftsZ2 cloned into pHVID7 | This study |
| pHVID62 | P.tnaA*_FtsZ2_Glink_$^{Hv}$YPet | NdeI-*ftsZ2*-BamHI from pIDJL40-ftsZ2 cloned into pHVID8 | This study |
| pHVID63 | P.tnaA*_FtsZ2_Glink_$^{Hv}$mTq-2 | NdeI-*ftsZ2*-BamHI from pIDJL40-ftsZ2 cloned into pHVID9 | This study |
| pHVID64 | P.tnaA*_FtsZ2_E_GFP | NdeI-*ftsZ2*-BamHI from pIDJL40-ftsZ2 cloned into pHVID10 | This study |
| pHVID65 | P.tnaA*_FtsZ2_E_$^{Hv}$mNG | NdeI-*ftsZ2*-BamHI from pIDJL40-ftsZ2 cloned into pHVID11 | This study |
| pHVID66 | P.tnaA*_FtsZ2_E_$^{Hv}$mCh | NdeI-*ftsZ2*-BamHI from pIDJL40-ftsZ2 cloned into pHVID12 | This study |
| pHVID67 | P.tnaA*_FtsZ2_E_$^{Hv}$mScar-I | NdeI-*ftsZ2*-BamHI from pIDJL40-ftsZ2 cloned into pHVID13 | This study |
| pHVID68 | P.tnaA*_FtsZ2_E_$^{Hv}$YPet | NdeI-*ftsZ2*-BamHI from pIDJL40-ftsZ2 cloned into pHVID14 | This study |
| pHVID69 | P.tnaA*_FtsZ2_E_$^{Hv}$mTq-2 | NdeI-*ftsZ2*-BamHI from pIDJL40-ftsZ2 cloned into pHVID15 | This study |
| pHVID70 | P.tnaA*_FtsZ2_EG_GFP | NdeI-*ftsZ2*-BamHI from pIDJL40-ftsZ2 cloned into pHVID16 | This study |
| pHVID71 | P.tnaA*_FtsZ2_EG_$^{Hv}$mNG | NdeI-*ftsZ2*-BamHI from pIDJL40-ftsZ2 cloned into pHVID17 | This study |
| pHVID72 | P.tnaA*_FtsZ2_EG_$^{Hv}$mCh | NdeI-*ftsZ2*-BamHI from pIDJL40-ftsZ2 cloned into pHVID18 | This study |
| pHVID73 | P.tnaA*_FtsZ2_EG_$^{Hv}$mScar-I | NdeI-*ftsZ2*-BamHI from pIDJL40-ftsZ2 cloned into pHVID19 | This study |
| pHVID74 | P.tnaA*_FtsZ2_EG_$^{Hv}$YPet | NdeI-*ftsZ2*-BamHI from pIDJL40-ftsZ2 cloned into pHVID20 | This study |
| pHVID75 | P.tnaA*_FtsZ2_EG_$^{Hv}$mTq-2 | NdeI-*ftsZ2*-BamHI from pIDJL40-ftsZ2 cloned into pHVID21 | This study |
| pHVID76 | P.tnaA*_GFP_Glink_FtsZ2 | PCR cloned into pHVID22 EcoRI/NheI<br>SI115_EcoRI TATGGAATTCCAGGATATCGTTCGCGAGGCG pHVID22<br>SI117_STOP-NheI<br>GTATGCTAGCTTACCGGATGACGTCGAGACCGTTGTTCTTCTCGACTTCGTCCCGGCCGC<br>Contains silent mutation to remove internal NotI site in *ftsZ2* (C1161G) | This study |
| pHVID77 | P.tnaA*_$^{Hv}$mNG_Glink_FtsZ2 | *ftsZ2* PCR fragment cloned into pHVID23 | This study |
| pHVID78 | P.tnaA*_$^{Hv}$mCh_Glink_FtsZ2 | *ftsZ2* PCR fragment cloned into pHVID24 | This study |
| pHVID79 | P.tnaA*_$^{Hv}$mScar-I_Glink_FtsZ2 | *ftsZ2* PCR fragment cloned into pHVID25 | This study |
| pHVID80 | P.tnaA*_$^{Hv}$YPet_Glink_FtsZ2 | *ftsZ2* PCR fragment cloned into pHVID26 | This study |
| pHVID81 | P.tnaA*_$^{Hv}$mTq-2_Glink_FtsZ2 | *ftsZ2* PCR fragment cloned into pHVID27 | This study |
| pHVID82 | P.tnaA*_GFP-E_FtsZ2 | *ftsZ2* PCR fragment cloned into pHVID28 | This study |
| pHVID83 | P.tnaA*_$^{Hv}$mNG_E_FtsZ2 | *ftsZ2* PCR fragment cloned into pHVID29 | This study |
| pHVID84 | P.tnaA*_$^{Hv}$mCh_E_FtsZ2 | *ftsZ2* PCR fragment cloned into pHVID30 | This study |
| pHVID85 | P.tnaA*_$^{Hv}$mScar-I-E_FtsZ2 | *ftsZ2* PCR fragment cloned into pHVID31 | This study |

| | | | |
|---|---|---|---|
| pHVID86 | P.tnaA*_$^{Hv}$YPet_E_FtsZ2 | *ftsZ2* PCR fragment cloned into pHVID32 | This study |
| pHVID87 | P.tnaA*_$^{Hv}$mTq-2_E_FtsZ2 | *ftsZ2* PCR fragment cloned into pHVID33 | This study |
| pHVID88 | P.tnaA*_GFP_EG_FtsZ2 | *ftsZ2* PCR fragment cloned into pHVID34 | This study |
| pHVID89 | P.tnaA*_$^{Hv}$mNG_EG_FtsZ2 | *ftsZ2* PCR fragment cloned into pHVID35 | This study |
| pHVID90 | P.tnaA*_$^{Hv}$mCh_EG_FtsZ2 | *ftsZ2* PCR fragment cloned into pHVID36 | This study |
| pHVID91 | P.tnaA*_$^{Hv}$mScar-I-EG_FtsZ2 | *ftsZ2* PCR fragment cloned into pHVID37 | This study |
| pHVID92 | P.tnaA*_$^{Hv}$YPet_EG_FtsZ2 | *ftsZ2* PCR fragment cloned into pHVID38 | This study |
| pHVID93 | P.tnaA*_$^{Hv}$mTq-2_EG_FtsZ2 | *ftsZ2* PCR fragment cloned into pHVID39 | This study |
| pHVID127 | P.tnaA*_FtsZ1_EG_$^{Hv}$mTq-2::$^{Hv}$mCh_EG_FtsZ2 | | This study |
| pHVID128 | P.tnaA*_$^{Hv}$mCh_EG_FtsZ2:: Fts[illegible] | | This study |
| pHVID129 | P.tnaA*_FtsZ1_EG_$^{Hv}$mTq-2::$^{Hv}$YPet_EG_FtsZ2 | | This study |
| pHVID130 | P.tnaA*_CetZ1_G_GFP | NdeI-*cetZ1*-BamHI from pIDJL40-*cetZ1* cloned into pHVID4 | This study |
| pHVID131 | P.tnaA*_CetZ1_G_$^{Hv}$mNG | NdeI-*cetZ1*-BamHI from pIDJL40-*cetZ1* cloned into pHVID5 | This study |
| pHVID132 | P.tnaA*_CetZ1_G_$^{Hv}$mCh | NdeI-*cetZ1*-BamHI from pIDJL40-*cetZ1* cloned into pHVID6 | This study |
| pHVID133 | P.tnaA*_CetZ1_G_$^{Hv}$mScar-I | NdeI-*cetZ1*-BamHI from pIDJL40-*cetZ1* cloned into pHVID7 | This study |
| pHVID134 | P.tnaA*_CetZ1_G_$^{Hv}$YPet | NdeI-*cetZ1*-BamHI from pIDJL40-*cetZ1* cloned into pHVID8 | This study |
| pHVID135 | P.tnaA*_CetZ1_G_$^{Hv}$mTq-2 | NdeI-*cetZ1*-BamHI from pIDJL40-*cetZ1* cloned into pHVID9 | This study |
| pHVID136 | P.tnaA*_CetZ1_E_GFP | NdeI-*cetZ1*-BamHI from pIDJL40-*cetZ1* cloned into pHVID10 | This study |
| pHVID137 | P.tnaA*_CetZ1_E_$^{Hv}$mNG | NdeI-*cetZ1*-BamHI from pIDJL40-*cetZ1* cloned into pHVID11 | This study |
| pHVID138 | P.tnaA*_CetZ1_E_$^{Hv}$mCh | NdeI-*cetZ1*-BamHI from pIDJL40-*cetZ1* cloned into pHVID12 | This study |
| pHVID139 | P.tnaA*_CetZ1_E_$^{Hv}$mScar-I | NdeI-*cetZ1*-BamHI from pIDJL40-*cetZ1* cloned into pHVID13 | This study |
| pHVID140 | P.tnaA*_CetZ1_E_$^{Hv}$YPet | NdeI-*cetZ1*-BamHI from pIDJL40-*cetZ1* cloned into pHVID14 | This study |
| pHVID141 | P.tnaA*_CetZ1_E_$^{Hv}$mTq-2 | NdeI-*cetZ1*-BamHI from pIDJL40-*cetZ1* cloned into pHVID15 | This study |
| pHVID142 | P.tnaA*_CetZ1_EG_GFP | NdeI-*cetZ1*-BamHI from pIDJL40-*cetZ1* cloned into pHVID16 | This study |
| pHVID143 | P.tnaA*_CetZ1_EG_$^{Hv}$mNG | NdeI-*cetZ1*-BamHI from pIDJL40-*cetZ1* cloned into pHVID17 | This study |
| pHVID144 | P.tnaA*_CetZ1_EG_$^{Hv}$mCh | NdeI-*cetZ1*-BamHI from pIDJL40-*cetZ1* cloned into pHVID18 | This study |
| pHVID145 | P.tnaA*_CetZ1_EG_$^{Hv}$mScar-I | NdeI-*cetZ1*-BamHI from pIDJL40-*cetZ1* cloned into pHVID19 | This study |
| pHVID146 | P.tnaA*_CetZ1_EG_$^{Hv}$YPet | NdeI-*cetZ1*-BamHI from pIDJL40-*cetZ1* cloned into pHVID20 | This study |
| pHVID147 | P.tnaA*_CetZ1_EG_$^{Hv}$mTq-2 | NdeI-*cetZ1*-BamHI from pIDJL40-*cetZ1* cloned into pHVID21 | This study |

*All of these plasmids are derivatives from a pTA962 backbone. P.tnaA* = modified p.tnaA promoter, G = fexible G linker (GGGGS)$_5$, E = rigid E linker (EAAAK)$_5$, EG = EG linker (EAAAK)$_1$(GGGGS)$_2$(EAAAK)$_2$, GFP = smRS-GFP, $^{Hv}$mNG = $^{Hv}$mNeonGreen, $^{Hv}$mCh = $^{Hv}$mCherry, $^{Hv}$mScar-I = $^{Hv}$mScarlett-I, $^{Hv}$mTq-2 = $^{Hv}$mTurquoise-2. Relevant restriction enzyme cutting sites for cloning are in bold, mutations are boxed.

**Supplementary Table 2.** DNA Sequences of promoters, linkers and genes encoding fluorescent proteins.

| Name | Abbreviation | DNA Sequence (5'-3') | Source |
|---|---|---|---|
| Modified *p.tnaA* | *p.tnaA** | CGTTCTCGTCGCGCTCTCGAAGCTGTTTCTCGCGCGCTCGCGTCCTCGAAAGTGACATCGCTCGACCGGTGGTCG<br>TCGGCGGTCGCTGAAGTCGGCTCGTGGCGAGAACGGAACAGCCGGCGACACCGATGCACACACCAGTCCACGA<br>GCGCCGAAAACCGGGCGTAGCCCCTCGATTTTCCGCCTGCCGATTACTTCACATTCGCGGACCTTCTAGACAT | This study |
| G linker | G | GGCGGCGGGGGCTCGGGGGGCGGCGGGTCGGGCGGCGGGGGGTCGGGCGGCGGGGGCAGCGGGGGCGGC<br>GGCTCC | This study |
| E linker | E | GAGGCCGCGGCCAAAGAAGCCGCCGCCAAGGAAGCGGCGGCCAAGGAGGCCGCCGCGAAAGAGGCGGCGGC<br>GAAG | This study |
| EG linker | EG | GAGGCCGCGGCCAAAGGGGGCGGCGGGTCGGGCGGCGGGGGGTCGGAGGCCGCCGCGAAAGAGGCGGCGG<br>CGAAG | This study |
| Soluble-modified Red-Shifted Green Fluorescent Protein | smRS-GFP | ATGAGTAAAGGAGAAGAACTTTTCACTGGAGTTGTCCCAATTCTTGTTGAATTAGATGGTGATGTTAATGGGCAC<br>AAATTTTCTGTCAGTGGAGAGGGTGAAGGTGATGCAACATACGGAAAACTTACCCTTAAATTTATTTGCACTACT<br>GGAAAACTACCTGTTCCATGGCCAACACTTGTCACTACTTTCACTTATGGTGTTCAATGCTTTTCAAGATACCCAG<br>ATCACATGAAGCGGCACGACTTCTTCAAGAGCGCCATGCCTGAGGGATACGTGCAGGAGAGGACCATCTCTTTC<br>AAGGACGACGGGAACTACAAGACACGTGCTGAAGTCAAGTTTGAGGGAGACACCCTCGTCAACAGGATCGAGC<br>TTAAGGGAATCGATTTCAAGGAGGACGGAAACATCCTCGGCCACAAGTTGGAATACAACTACAACTCCCACAAC<br>GTATACATCACGGCAGACAAACAAAAGAATGGAATCAAAGCTAACTTCAAAATTAGACACAACATTGAAGATGG<br>AAGCGTTCAACTAGCAGACCATTATCAACAAAATACTCCAATTGGCGATGGCCCTGTCCTTTTACCAGACAACCAT<br>TACCTGTCCACACAATCTGCCCTTTCGAAAGATCCCAACGAAAAGAGAGACCACATGGTCCTTCTTGAGTTTGTAA<br>CAGCTGCTGGGATTACACATGGCATGGATGAACTATACAAATAA | [5] |
| mCherry | mCh | GTGAGCAAGGGCGAGGAGGATAACATGGCTATCATTAAAGAGTTCATGCGCTTCAAAGTTCACATGGAGGGTTC<br>TGTTAACGGTCACGAGTTCGAGATCGAAGGCGAAGGCGAGGGCCGTCCGTATGAAGGCACCCAGACCGCCAAA<br>CTGAAAGTGACTAAAGGCGGCCCGCTGCCTTTTGCGTGGGACATCCTGAGCCCGCAATTTATGTACGGTTCTAAA<br>GCTTATGTTAAACACCCAGCGGATATCCCGGACTATCTGAAGCTGTCTTTTCCGGAAGGTTTCAAGTGGGAACGC<br>GTAATGAATTTTGAAGATGGTGGTGTCGTGACCGTCACTCAGGACTCCTCCCTGCAGGATGGCGAGTTCATCTAT<br>AAAGTTAAACTGCGTGGTACTAATTTTCCATCTGATGGCCCGGTGATGCAGAAGAAGACGATGGGTTGGGAGGC<br>GTCTAGCGAACGCATGTATCCGGAAGATGGTGCGCTGAAAGGCGAAATTAAACAGCGCCTGAAACTGAAAGAT<br>GGCGGCCATTATGACGCTGAAGTGAAAACCACGTACAAAGCCAAGAAACCTGTGCAGCTCCCTGGCGCGTACAA<br>TGTGAATATTAAACTGGACATCACCTCTCATAATGAAGATTATACGATCGTAGAGCAATATGAGCGCGCGGAGG<br>GTCGTCATTCTACCGGTGGCATGGACGAGCTGTACAAGTAA | [5] |
| $^{Hv}$mNeonGreen | $^{Hv}$mNG | GTCTCGAAGGGCGAGGAAGACAACATGGCCTCGCTCCCCGCCACCCACGAGCTCCACATCTTCGGCTCGATCAA<br>CGGCGTCGACTTCGACATGGTCGGCCAGGGCACCGGCAACCCCAACGACGGCTACGAGGAGCTCAACCTCAAGT<br>CGACCAAGGGCGACCTGCAGTTCTCGCCGTGGATTCTCGTCCCCCACATCGGCTACGGCTTCCACCAGTACCTGC<br>CGTACCCGGACGGCATGTCGCCCTTCCAGGCGGCCATGGTCGACGGCTCGGGCTACCAGGTCCACCGCACCATG<br>CAGTTCGAGGACGGCGCCTCGCTCACGGTCAACTACCGCTACACCTACGAGGGCTCGCACATCAAGGGCGAGGC<br>CCAGGTCAAGGGCACCGGCTTCCCGGCGGACGGCCCGGTCATGACGAACTCGCTCACCGCCGCGGACTGGTGC<br>CGCTCGAAGAAGACCTACCCCAACGACAAGACGATCATCTCGACCTTCAAGTGGTCGTACACGACCGGCAACGG<br>CAAGCGCTACCGCTCGACCGCCCGCACGACCTACACCTTCGCGAAGCCGATGGCCGCGAACTACCTCAAGAACC<br>AGCCCATGTACGTCTTCCGCAAGACGGAGCTCAAGCACTCGAAGACCGAGCTCAACTTCAAGGAGTGGCAGAAG<br>GCGTTCACGGACGTCATGGGCATGGACGAGCTCTACAAGTGA | This study |
| $^{Hv}$mCherry | $^{Hv}$mCh | GTCTCGAAGGGCGAGGAGGACAACATGGCGATCATCAAGGAGTTCATGCGCTTCAAGGTCCACATGGAGGGCT<br>CGGTCAACGGCCACGAGTTCGAGATCGAGGGCGAGGGCGAGGGCCGCCCGTACGAGGGCACGCAGACCGCCA<br>AGCTCAAGGTCACGAAGGGCGGCCCGCTCCCCTTCGCGTGGGACATCCTCTCGCCGCAGTTCATGTACGGCTCG<br>AAGGCCTACGTCAAGCACCCGGCGGACATCCCCGACTACCTCAAGCTCTCGTTCCCCGAGGGCTTCAAGTGGGA<br>GCGCGTCATGAACTTCGAGGACGGCGGCGTCGTCACGGTCACCCAGGACTCGTCGCTCCAGGACGGCGAGTTCA<br>TCTACAAGGTCAAGCTCCGCGGCACGAACTTCCCGTCGGACGGCCCCGTCATGCAGAAGAAGACCATGGGCTGG<br>GAGGCCTCGTCGGAGCGCATGTACCCCGAGGACGGCGCGCTCAAGGGCGAGATCAAGCAGCGCCTCAAGCTCA<br>AGGACGGCGGCCACTACGACGCCGAGGTCAAGACGACCTACAAGGCGAAGAAGCCGGTCCAGCTCCCCGGCGC<br>CTACAACGTCAACATCAAGCTCGACATCACGTCGCACAACGAGGACTACACCATCGTCGAGCAGTACGAGCGCG<br>CCGAGGGCCGCCACTCCACGGGCGGCATGGACGAGCTCTACAAGTAA | This study |
| $^{Hv}$mScarlet-I | $^{Hv}$mScar | GTCTCGAAGGGCGAGGCCGTCATCAAGGAGTTCATGCGCTTCAAGGTCCACATGGAGGGCTCGATGAACGGCC<br>ACGAGTTCGAGATCGAGGGCGAGGGCGAGGGCCGCCCGTACGAGGGCACGCAGACCGCGAAGCTCAAGGTCA<br>CCAAGGGCGGCCCCCTCCCCTTCTCGTGGGACATCCTCTCGCCGCAGTTCATGTACGGCTCGCGCGCCTTCATCA<br>AGCACCCGGCGGACATCCCCGACTACTACAAGCAGTCGTTCCCCGAGGGCTTCAAGTGGGAGCGCGTCATGAAC<br>TTCGAGGACGGTGGCGCCGTCACCGTCACGCAGGACACGTCGCTCGAGGACGGCACCCTCATCTACAAGGTCAA<br>GCTCCGCGGCACGAACTTCCCGCCCGACGGCCCCGTCATGCAGAAGAAGACGATGGGCTGGGAGGCCTCGACG<br>GAGCGCCTCTACCCCGAGGACGGCGTCCTCAAGGGCGACATCAAGATGGCCCTCCGCCTCAAGGACGGCGGCC<br>GGTACCTCGCCGACTTCAAGACGACCTACAAGGCGAAGAAGCCCGTCCAGATGCCGGGCGCGTACAACGTCGAC<br>CGCAAGCTCGACATCACGTCGCACAACGAGGACTACACCGTCGTCGAGCAGTACGAGCGCTCGGAGGGCCGCC<br>ACTCGACGGGCGGCATGGACGAGCTCTACAAGTGA | This study |
| $^{Hv}$YPet | $^{Hv}$YPet | TCGAAGGGCGAGGAGCTCTTCACGGGCGTCGTCCCCATCCTCGTCGAGCTCGACGGCGACGTCAACGGCCACAA<br>GTTCTCGGTCTCGGGCGAGGGCGAGGGCGACGCCACGTACGGCAAGCTCACCCTCAAGCTCCTCTGCACCACGG<br>GCAAGCTCCCCGTCCCCTGGCCCACCCTCGTCACCACGCTCGGCTACGGCGTCCAGTGCTTCGCCCGCTACCCCG<br>ACCACATGAAGCAGCACGACTTCTTCAAGTCGGCCATGCCCGAGGGCTACGTCCAGGAGCGCACCATCTTCTTCA<br>AGGACGACGGCAACTACAAGACGCGCGCCGAGGTCAAGTTCGAGGGCGACACCCTCGTCAACCGCATCGAGCT<br>CAAGGGCATCGACTTCAAGGAAGACGGCAACATCCTCGGCCACAAGCTCGAGTACAACTACAACTCGCACAACG<br>TCTACATCACGGCCGACAAGCAGAAGAACGGCATCAAGGCGAACTTCAAGATCCGCCACAACATCGAGGACGG | This study |

| | | | |
|---|---|---|---|
| | | CGGCGTCCAGCTCGCCGACCACTACCAGCAGAACACGCCGATCGGCGACGGCCCGGTCCTCCTCCCGGACAACC<br>ACTACCTCTCGTACCAGTCGGCGCTCTTCAAGGACCCGAACGAGAAGCGCGACCACATGGTCCTCCTCGAGTTCC<br>TCACGGCCGCGGGCATCACGGAGGGCATGAACGAGCTCTACAAGTGA | |
| $^{Hv}$mTurquoise-2 | $^{Hv}$mTq-2 | GTCTCGAAGGGCGAGGAGCTCTTCACGGGCGTCGTCCCCATCCTCGTCGAGCTCGACGGCGACGTCAACGGCCA<br>CAAGTTCTCGGTCTCGGGCGAGGGCGAGGGCGACGCCACGTACGGCAAGCTCACCCTCAAGTTCATCTGCACGA<br>CGGGCAAGCTCCCCGTCCCGTGGCCCACCCTCGTCACCACGCTCTCGTGGGGCGTCCAGTGCTTCGCCCGCTACC<br>CCGACCACATGAAGCAGCACGACTTCTTCAAGTCGGCCATGCCCGAGGGCTACGTCCAGGAGCGCACCATCTTCT<br>TCAAGGACGACGGCAACTACAAGACGCGCGCCGAGGTCAAGTTCGAGGGCGACACCCTCGTCAACCGCATCGA<br>GCTCAAGGGCATCGACTTCAAGGAAGACGGCAACATCCTCGGCCACAAGCTCGAGTACAACTACTTCTCGGACA<br>ACGTCTACATCACCGCCGACAAGCAGAAGAACGGCATCAAGGCGAACTTCAAGATCCGCCACAACATCGAGGAC<br>GGCGGCGTCCAGCTCGCGGACCACTACCAGCAGAACACGCCCATCGGCGACGGCCCGGTCCTCCTCCCGGACAA<br>CCACTACCTCTCGACCCAGTCGAAGCTCTCGAAGGACCCGAACGAGAAGCGCGACCACATGGTCCTCCTCGAGTT<br>CGTCACGGCCGCGGGCATCACCCTCGGCATGGACGAGCTCTACAAGTGA | This study |
